# Vertical rooting caused by enhanced functional allele of *qSOR1* improves rice yield under drought stress

**DOI:** 10.1101/2025.06.07.658467

**Authors:** Tsubasa Kawai, Yuka Kitomi, Ryo Kuroda, Ryu Nakata, Momoha Iba, Fumiyuki Soma, Shota Teramoto, Toshimasa Yamazaki, Kazuhiko Sugimoto, Yusaku Uga

## Abstract

Drought considerably affects crop productivity, and its severity is being intensified by climate change. Therefore, enhancing drought resistance is a crucial priority in crop breeding for ensuring sustainable agriculture. The root system architecture (RSA) influences the efficiency of water acquisition from land; therefore, a deep RSA is advantageous for avoiding drought stress. Here, we demonstrated that deeper RSA promoted by the *qSOR1*-v mutant allele (an enhanced functional allele of the *quantitative trait locus for SOIL SURFACE ROOTING 1*) significantly improves rice yield under drought when compared to the deep RSA achieved through the functional *qSOR1* allele that originated from natural variation. The *qSOR1*-v mutant exhibited stronger root gravitropism than the wild type. This was characterized by a more pronounced polarization of auxin on the lower side during root curvature, leading to a robust vertical rooting phenotype that was consistently expressed across different soil-water environments. Additionally, the *qSOR1*-v mutation site was well conserved among angiosperm orthologs, and the corresponding mutation in *LZY3* of *Arabidopsis* (*qSOR1* ortholog) resulted in a steeper root growth angle. The rice introgression line, which was substituted from the functional *qSOR1* allele to *qSOR1*-v through marker-assisted selection, showed vertical rooting, resulting in increased grain yield in an upland field under drought stress. No yield penalty was observed for this line under well-watered upland conditions than the original variety. These findings highlight the potential of *qSOR1*-v and corresponding mutations in angiosperm orthologs to promote vertical rooting across plant species, which can help sustain crop yields in drought-prone areas.

**Significance Statement:** Genetic modification of the root system architecture in crops represents a viable strategy for the development of climate-resilient crops. This study identified the *qSOR1*-v allele that consistently demonstrates a vertical rooting phenotype in rice across diverse growth conditions, from dry to wet. The preservation of the qSOR1 and LZY3 orthologs in angiosperms provides opportunities for the development of genotypes characterized by vertical rooting. The introgression of the *qSOR1*-v allele enhanced drought resistance under upland conditions. These findings underscore the potential of these genetic modifications to improve crop resilience in an era characterized by water scarcity.

## Introduction

Drought is a major constraint on agricultural productivity, limiting vegetative and reproductive processes and ultimately leading to yield loss (1, 2). Owing to decreased precipitation and changes in rainfall patterns driven by climate change, crop production is increasingly exposed to drought stress, posing a serious threat to global food security (3). Improving drought resistance is an urgent challenge for sustainable crop production.

Rice is cultivated under various hydrological conditions, such as irrigated lowlands, rainfed lowlands, and rainfed uplands. The rainfed rice area covers approximately half of the total cultivated area but contributes only approximately 20% of the total rice production (4). Rainfed rice production tends to produce lower average yields owing to limited and unpredictable water availability, low inputs, and suboptimal management practices, and is more vulnerable to climate change (5, 6). In addition to improved management practices, breeding drought-resistant rice is a key strategy for mitigating the effects of drought by targeting traits related to drought escape, avoidance, and tolerance (7).

Roots are essential organs for the uptake of water and nutrients from the soil. Root system architecture (RSA), which is determined by the number of roots formed, root elongation, and root growth angle (RGA), considerably affects the efficiency of water and nutrient acquisition in the soil (8). RSA ideotypes differ across soil environments, particularly in those with heterogeneous water and nutrient distributions (9, 10). Several studies have demonstrated that deep rooting is valuable for drought avoidance in rainfed fields. Under drought conditions, greater root distribution at depth is associated with enhanced water uptake (11, 12), resulting in reduced yield loss among rice cultivars (13). Among rice cultivars, a greater ability to increase root length density at depth in response to soil moisture availability is associated with higher drought resistance (14, 15). Similar to rice, the contribution of deep rooting to increased drought resistance has been reported in wheat (16, 17), maize (18), pearl millet (19), common bean (20), and chicory (21). However, to date, only a few genes that confer a deep rooting phenotype have been identified. In rice, *DEEPER ROOTING1* (*DRO1*) was first identified as a quantitative trait locus (QTL) controlling RGA in a cross between the lowland cultivar IR64 and the upland cultivar Kinandang Patong (KP) (22). The introduction of the functional allele of *DRO1* from KP to IR64 conferred deep rooting, resulting in improved yields under upland drought conditions (23, 24). *Rice morphology determinant* (*RMD*), which encodes a rice actin-binding protein, regulates RGA in response to phosphate, and its mutation results in deep root growth (25). In maize, genome-wide association mapping identified *ZmCIPK15* as a gene that regulates RGA, and its mutation resulted in increased RGA levels (26). In barley, *ENHANCED GRAVITROPISM 1* (*EGT1*) was identified as the causative gene of a mutant exhibiting deeper root growth, and mutations in orthologous *EGT1* genes in wheat resulted in steeper RGA (27). These genetic findings are valuable resources for improving deep rooting in crops. However, the extent of RGA modification may vary depending on the gene/allele and genetic background. Their potential to enhance crop productivity, particularly under drought stress, remains poorly understood.

The regulation of RGA is associated with a gravitropic response in the root tips. Root gravitropism involves the following steps: gravity sensing and signaling in columella cells, intercellular signal transduction via auxins, and asymmetric organ growth in the root elongation zone (28). *LAZY1*-family genes, a subgroup of the IGT family (29), play a major role in gravitropism. *LAZY1* (*LA1*) was first identified in rice as the causative gene of a mutant exhibiting prostrate growth habit in the shoots (30, 31). A homology search of the closest *Arabidopsis* ortholog, *AtLAZY1/LAZY1-like 1* (*LZY1*; the nomenclature of *Arabidopsis LAZY1*-family genes is based on the classification by Taniguchi *et al*. [32]), identified five additional homologs in *Arabidopsis* (33). Several studies have revealed that *LZY1*, *LZY2* (*AtLAZY2*/*AtNGR1*/*AtDRO3*), and *LZY3* (*AtLAZY4*/*AtNGR2*/*AtDRO1*) are involved in shoot gravitropism, whereas *LZY2*, *LZY3*, and *LZY4* (*AtLAZY3*/*AtNGR3*/*AtDRO2*) are involved in root gravitropism (32, 34–36). Recent studies revealed a gravity-sensing mechanism involving LZY3 in columella cells (37, 38). Following gravitational stimulation, LZY3 is translocated from the amyloplast to the lower plasma membrane. This localization promotes the recruitment of RCC1-like domain (RLD) proteins to the same region, subsequently modulating the polar localization of PIN auxin efflux carriers (39). The unequal distribution of auxins between the upper and lower sides of gravity-stimulated roots induces differential cell elongation, resulting in root curvature toward a new gravity vector (40). In rice, the closest ortholog of *LZY3* is the *quantitative trait locus for SOIL SURFACE ROOTING 1* (*qSOR1*), a *DRO1* homolog that was identified as a QTL controlling RGA in a cross between the lowland cultivars Sasanishiki and Gemdja Beton (GB) (41, 42). *DRO1* and *qSOR1* additively modulate RSA; combinations of functional and non-functional alleles result in continuous variation from ultra-shallow to deep rooting (42). *LAZY1*-family genes have been identified, and their functions in RGA regulation have been investigated in other plant species, including *Medicago* (34), plum (35), wheat (43), barley (44), and cucumber (45), highlighting the significance of *LAZY1*-family genes for RSA modification across plant species.

In this study, allele mining of rice genes regulating RGA, *DRO1* and *qSOR1*, was conducted to identify new alleles for modulating RSA, in order to test whether further enhanced deeper root growth (namely vertical rooting) confers improved drought resistance in rice. The newly identified *qSOR1*-v allele was introgressed into two different genetic backgrounds, combined with functional and non-functional *DRO1*, to evaluate the effect of the allelic combination on RSA and drought resistance. The well-conserved *qSOR1*-v mutation site in orthologs, along with the resulting steeper RGA caused by the corresponding mutation in *LZY3* of *Arabidopsis*, indicates the potential application of these mutations in modulating RSA across plant species.

## Results

### Isolation of *qSOR1* vertical rooting mutants by TILLING

To obtain genetic resources to modulate RSA in rice, we screened *DRO1* and *qSOR1* mutant alleles in the background of a lowland rice cultivar ‘Koshihikari (KSH)’ using the targeting-induced local lesions (TILLING) system (46). Although no allelic variants were found in *DRO1*, three allelic variants with non-synonymous SNPs were identified in *qSOR1* (Fig. 1A). Root phenotypes were analyzed using the basket method (23, 24). The two mutants, 0951M (P140S) and 2792M (L141F), showed a steeper RGA, and the other mutant, 0909M (R204C), exhibited a shallower RGA than the wild type (WT) (Fig. 1B, C). The ratio of deep rooting (≥ 50°) was higher in 0951M and 2792M and lower in 0909M than that in the WT (Fig. 1D). These results indicated that the 0951M and 2792M lines exhibited vertical rooting. The root phenotypes of the backcrossing populations of 2792M were segregated in a 1:2:1 WT:intermediate:mutant ratio, suggesting that the mutant alleles were semi-dominant. Herein, the two mutants (alleles) with vertical rooting are referred to as *qSOR1*-v mutants (alleles).

**Figure 1.**
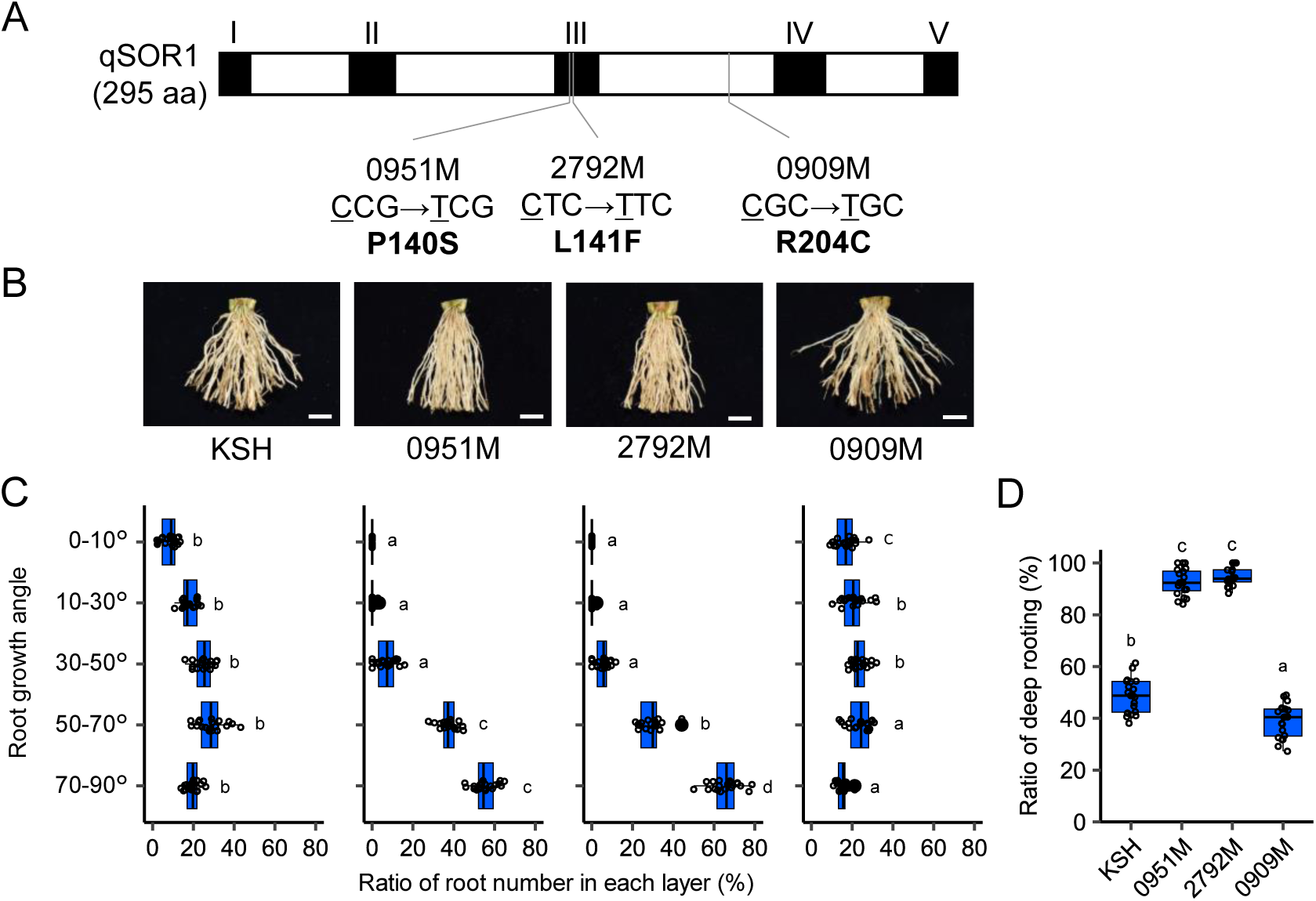
Characterization of *qSOR1* mutants. (A) Domain structure of qSOR1 protein and mutated sites in the three mutants. The black boxes represent regions I–V. (B) Images of basal parts of rice plants grown in the basket for 35 days. KSH, Koshihikari. Scale bars, 1 cm. (C, D) Ratio of root number (C) in each layer and (D) in the deeper layer (≥ 50°) to the total number in (B). Boxes represent the first quartile, median, and third quartile. Whiskers show the range of nonoutlier values. Filled circles indicate outlier values less than the first quartile or greater than the third quartile by 1.5 times the interquartile range. Different lowercase letters indicate significant differences among layers (*p* < 0.05, multiple-comparison Tukey’s test, *n* = 20 plants).

The *qSOR1* expression level in the root tips was slightly higher in 2792M than in the WT (approximately 1.5 times), while the levels in 0951M and 0909M were similar to that in the WT (*SI Appendix*, Fig. S1A). The mutated sites of *qSOR1* in the 0951M and 2792M lines are located in region III among the five conserved domains of qSOR1 and orthologous proteins (Fig. 1A; ref. 33). Three-dimensional (3D) modeling of qSOR1 proteins showed that R204 might affect the stability of the helix bundle through the formation of salt bridges with D17 and E266. R204C could inhibit its role, and P140S and L141F likely do not affect the protein structure (*SI Appendix*, Fig. S2). Both WT and mutated qSOR1 (L141F) were localized to the plasma membrane and plastids in rice protoplasts (*SI Appendix*, Fig. S3). These results indicate that mutations in *qSOR1*-v alleles may affect *qSOR1* gene function(s) other than its gene expression level, protein localization, and structure.

### Auxin redirection during the gravitropic response in the WT and *qSOR1* mutants

Because the phytohormone auxin plays a pivotal role in root gravitropism, we analyzed auxin redirection during the gravitropic response in the WT and *qSOR1* mutants. The degree of root curvature after rotation was higher in the 0951M and 2792M mutants and lower in the 0909M mutant than in the WT (Fig. 2A, B). The expression pattern of the auxin response gene *OsIAA20* (47) was analyzed during redirection in the elongation zone of the seminal root tip. After root rotation, *OsIAA20* expression was higher on the lower side than on the upper side of the rotated roots in the WT, and its ratio was notably higher in 0951M and 2792M (approximately 3 times versus approximately 1.5 times in the WT, Fig. 2C). In contrast, *OsIAA20* expression levels were not significantly different between the upper and lower sides in 0909M (Fig. 2C). The endogenous IAA level was higher in the lower side than in the upper side in 2792M, but not in WT (Fig. 2D). This suggests that *qSOR1*-v induces increased auxin accumulation on the lower side of the curvature, resulting in a higher gravitropic response in the roots.

**Figure 2.**
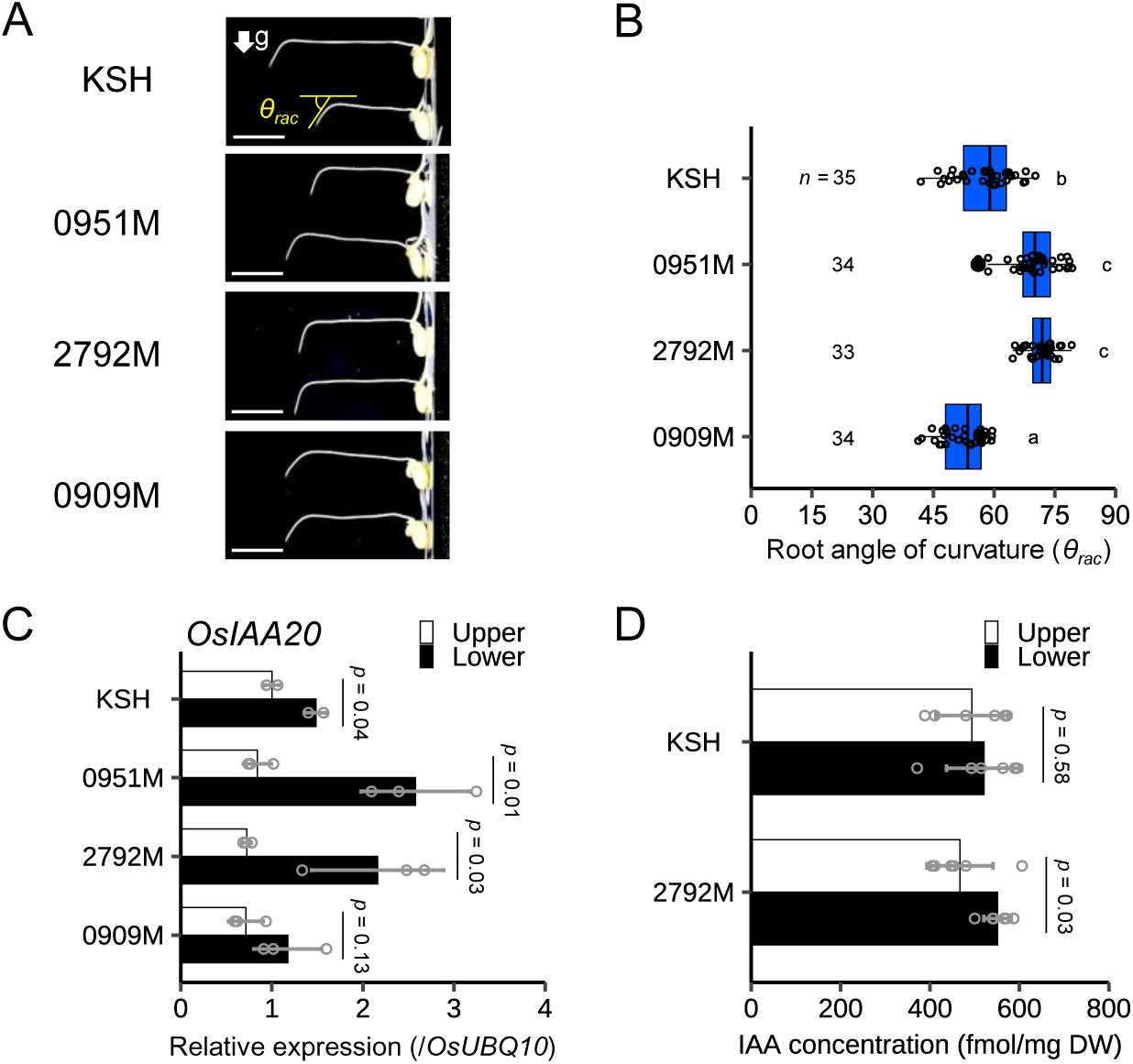
Root gravitropic response in *qSOR1* mutants. (A) Images of gravitropic curvature in the seminal roots after 4 h of 90° rotation. KSH, Koshihikari. *θ_rac_* is the root angle of curvature after the rotation. The arrow indicates the direction of the gravitational force. Scale bars, 1 cm. (B) Root angle of curvature in (A). Boxes represent the first quartile, median, and third quartile. Whiskers show the range of nonoutlier values. Filled circles indicate outlier values less than the first quartile or greater than the third quartile by 1.5 times the interquartile range. Different lowercase letters indicate significant differences among groups (*p* < 0.05, multiple-comparison Tukey’s test). (C, D) *OsIAA20* expressions (C) (*n* = 2 or 3 biological replicates) and endogenous IAA concentration (D) (*n* = 6 biological replicates) in the upper and lower parts of seminal root tips (0.5–2.5 cm from the root tip) after 1 h of the rotation. The expression of *OsIAA20* was normalized to that of *OsUBQ10*. Bar plots show mean values ± SD. *p* values are derived from a two-tailed Student’s *t*-test.

### Conserved effect of *qSOR1*-v mutation on RGA in rice and *Arabidopsis*

Phylogenetic analysis of DRO1 and LAZY1 clade proteins from the IGT family identified five subgroups in the DRO1 clade: qSOR1/DRL1, LZY2/3, DRO1-monot, DRO1-dicot, and DRL2 subgroups (*SI Appendix*, Fig. S4A). Alignment analysis revealed that the amino acid sequence of the qSOR1 region III, the mutated site of *qSOR1*-v, was highly conserved in the qSOR1/DRL1 and LZY2/3 subgroups but not in the other subgroups (Fig. 3A, *SI Appendix*, Fig. S4, S5). To analyze whether the point mutation in region III causes the vertical rooting phenotype in dicotyledonous plant species, we transformed the mutated *LZY3* in *Arabidopsis* into the *lzy2 lzy3* mutant with a shallower lateral root angle (*SI Appendix*, Fig. S1B, C; ref. 32). Transformants with the single-copy WT allele of *LZY3* into the *lzy2 lzy3* mutant, which was validated by expression analysis in individuals (*SI Appendix*, Fig. S1D), completely recovered the shallow rooting of the mutant (Fig. 3B, C). Transformants of the single-copy v-type mutated *LZY3* (*LZY3*-v, L131F) into the *lzy2 lzy3* mutant showed steeper lateral root angles in two independent lines (#2-7 and #8-3) (Fig. 3B, C, *SI Appendix*, Fig. S1D). This indicated the conserved roles of *qSOR1*-v and *LZY3*-v mutations in promoting vertical rooting in rice and *Arabidopsis*.

**Figure 3.**
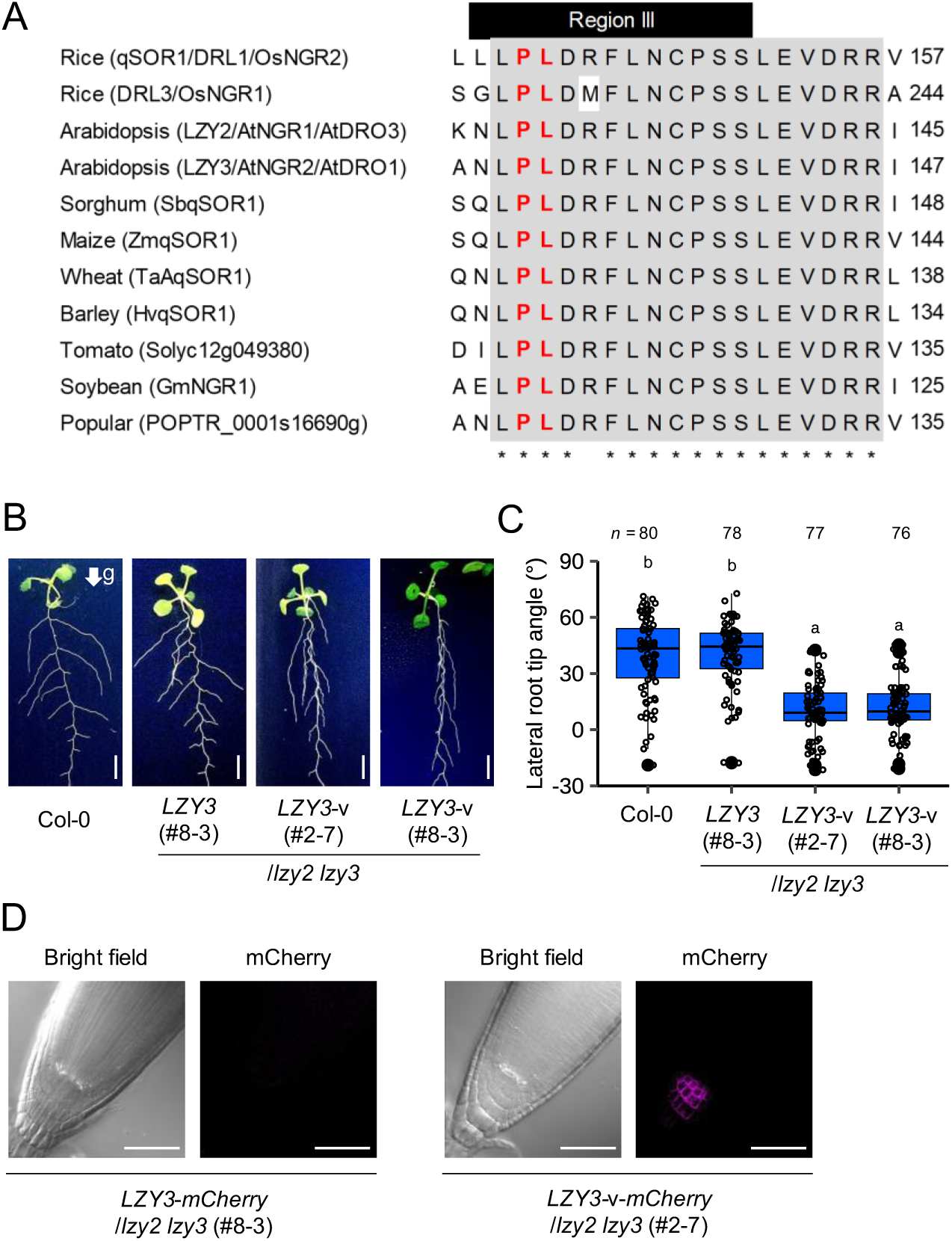
Comparative analysis of the vicinity of *qSOR1*-v mutation sites and the effects of the corresponding mutation in *Arabidopsis*. (A) Multiple sequence alignment of the vicinity of region III from qSOR1 orthologous proteins. Amino acid residues highlighted with a gray background and an asterisk are conserved. The two amino acid residues shown in red indicate mutation sites in the two *qSOR1*-v mutants (P140S in 0951M and L141F in 2792M). (B) Images of *Arabidopsis* plants of wild type (Col-0) and *lzy2 lzy3* transgenic lines harboring a single copy of either *LZY3* or *LZY3*-v (L131F). The arrow indicates the direction of the gravitational force. Scale bars, 5 mm. (C) Lateral root tip angle in (B). The angle of each root tip relative to the gravity vector was measured. Boxes represent the first quartile, median, and third quartile. Whiskers show the range of nonoutlier values. Filled circles indicate outlier values less than the first quartile or greater than the third quartile by 1.5 times the interquartile range. Different lowercase letters indicate significant differences among groups (*p* < 0.05, multiple-comparison Tukey’s test). (D) Expression of wild-type *LZY3* and *LZY3*-v fused to the mCherry fluorescent protein in *Arabidopsis* root tips. Scale bars, 30 μm.

To analyze the dose effect of *LZY3* and *LZY3*-v expressions, we obtained transformants with various *LZY3* expression levels and measured their root phenotypes. The transformant with 3.8 times higher *LZY3* expression (#1-3) exhibited a steeper root angle than that of the WT (*SI Appendix*, Fig. S6). However, the higher expression of *LZY3* (#6-2:10.6 times) and *LZY3*-v (#2-1-2:4.2 times, #4-1:6.1 times) resulted in shallower root angles (*SI Appendix*, Fig. S6). This suggests a dose-dependent effect of *LZY3* expression on root angle: moderate overexpression of WT *LZY3* promoted the deep rooting phenotype, whereas introducing single-copy *LZY3*-v enhanced vertical rooting the most.

Next, the expression of *LZY3* and *LZY3*-v was analyzed in the root tips of *Arabidopsis* in single-copy *LZY3*-*mCherry*/*lzy2 lzy3* (#8-3) and *LZY3*-v-*mCherry*/*lzy2 lzy3* (#2-7) transformants (Fig. 3B, C, *SI Appendix*, Fig. S1D). The LZY3-mCherry signal was barely detectable in root tips (Fig. 3D). However, LZY3-v-mCherry was detected in columella cells, a gravity-sensing organ (Fig. 3D). The application of the protein degradation inhibitor MG132 resulted in increased LZY3-mCherry signals in the root tips, including columella cells (*SI Appendix*, Fig. S7). These results indicated that v-type mutations affect LZY3 protein abundance or turnover rate.

### Combination of *qSOR1*-v with *DRO1* alleles expands rice RSA phenotypes

In our previous study, we developed three introgression lines (ILs) carrying *DRO1* and *qSOR1* alleles in an IR64 (*indica*) background (42). To investigate the effects of *qSOR1*-v in combination with *DRO1* on RSA modification, we developed two ILs harboring *qSOR1*-v with *DRO1*[KP] or *dro1*[IR64] in the IR64 background (*SI Appendix*, Table S1). Time-course image analysis using X-ray computed tomography (CT) for rice RSA development in the plant pot (48, 49) revealed that six combinations of *DRO1* and *qSOR1* exhibited varying root distributions in their RSAs (Fig. 4A). Root depth index (RDI) was highest in DRO1+qSOR1-v_PYL, followed by qSOR1-v_NIL, DRO1_NIL, DRO1+qsor1_PYL, IR64, and qsor1_NIL (Fig. 4B). The order of ILs was the same for RGA (Fig. 4C), which had significant positive correlations with the RDI after 21 days after sowing (DAS) (*SI Appendix*, Fig. S8C). Root number (RN) and total root length (TRL) showed similar trends, with a significant positive correlation with the RDI for TRL at 35 DAS (*SI Appendix*, Fig. S8A, B, E).

**Figure 4.**
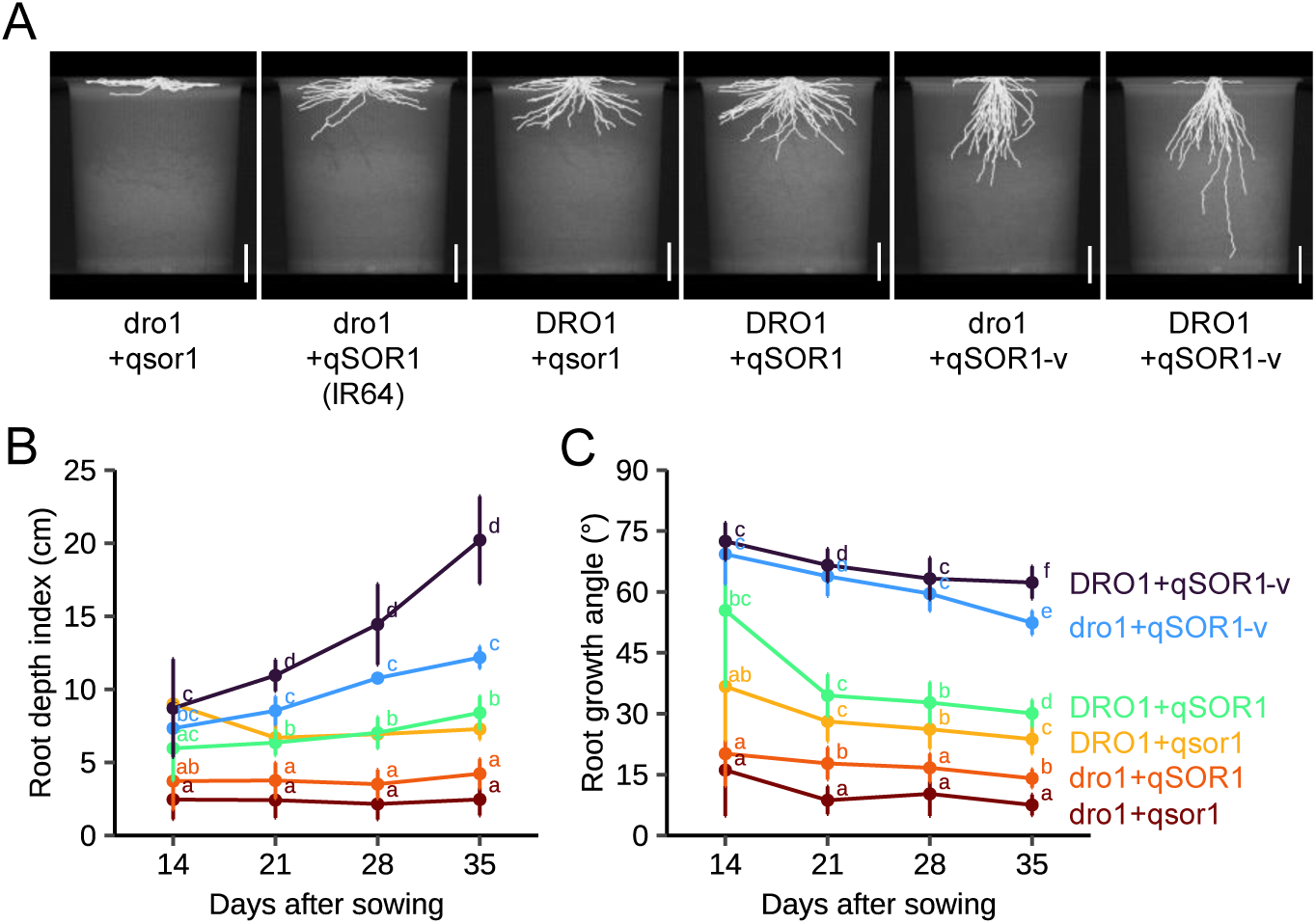
RSA analysis of IR64 and the five ILs with different *DRO1* and *qSOR1* allele combinations grown in a plant pot under a controlled chamber. (A) Images of RSA in 35-day-old rice plants visualized by X-ray CT and image processing. Scale bars, 5 cm. (B, C) Time-course analyses of (B) the root depth index and (C) root growth angle. Line charts show mean values ± SD (*n* = 6 biological replicates). Different lowercase letters indicate significant differences among groups (*p* < 0.05, multiple-comparison Tukey’s test).

To investigate the effects of *DRO1* and *qSOR1* alleles on RSA in different genetic backgrounds, we produced five ILs in a KSH (*japonica*) background (*SI Appendix*, Table S1, Fig. S9). In the KSH background, ILs with *qSOR1*-v showed significantly deep RSAs; however, the effect of *qsor1* derived from 0909M (cf. derived from GB in the IR64 background) was not significant for RDI and RGA (*SI Appendix*, Fig. S9B, C). Significant positive correlations with RDI were detected for RGA but not for RN or TRL (*SI Appendix*, Fig. S9F–H). These results suggest that the combination of *qSOR1*-v and *DRO1* contributes to the vertical rooting phenotype in different genotypes.

### Different genotypes exhibit varying relationships with RSA and yield in the paddy field

To verify the RSA differences in the *DRO1 qSOR1* ILs and their relationship with shoot growth and yield performance, RSA and yield components were evaluated in the five ILs and parents in the IR64 and KSH backgrounds in an irrigated paddy field. In the IR64 background, image analysis using X-ray CT for RSA in soil monoliths sampled from the paddy (50) revealed continuous variation in RGA due to different allele combinations of *DRO1* and *qSOR1* (Fig. 5A, B). The root diameter (RD) was slightly greater in ILs with *qSOR1*-v than in the other lines (*SI Appendix*, Fig. S10A). RN was lower in qsor1_NIL than in the other lines (*SI Appendix*, Fig. S10B). In the KSH background, ILs with *qSOR1*-v showed significantly deep RSAs with minor effects of *qsor1* (*SI Appendix*, Fig. S13A, B). *qSOR1*-v slightly increased RD, with no difference in RN among the six lines (*SI Appendix*, Fig. S13C, D). These observations verify the effect of *qSOR1*-v on deep RSA in both genotypes under paddy conditions.

**Figure 5.**
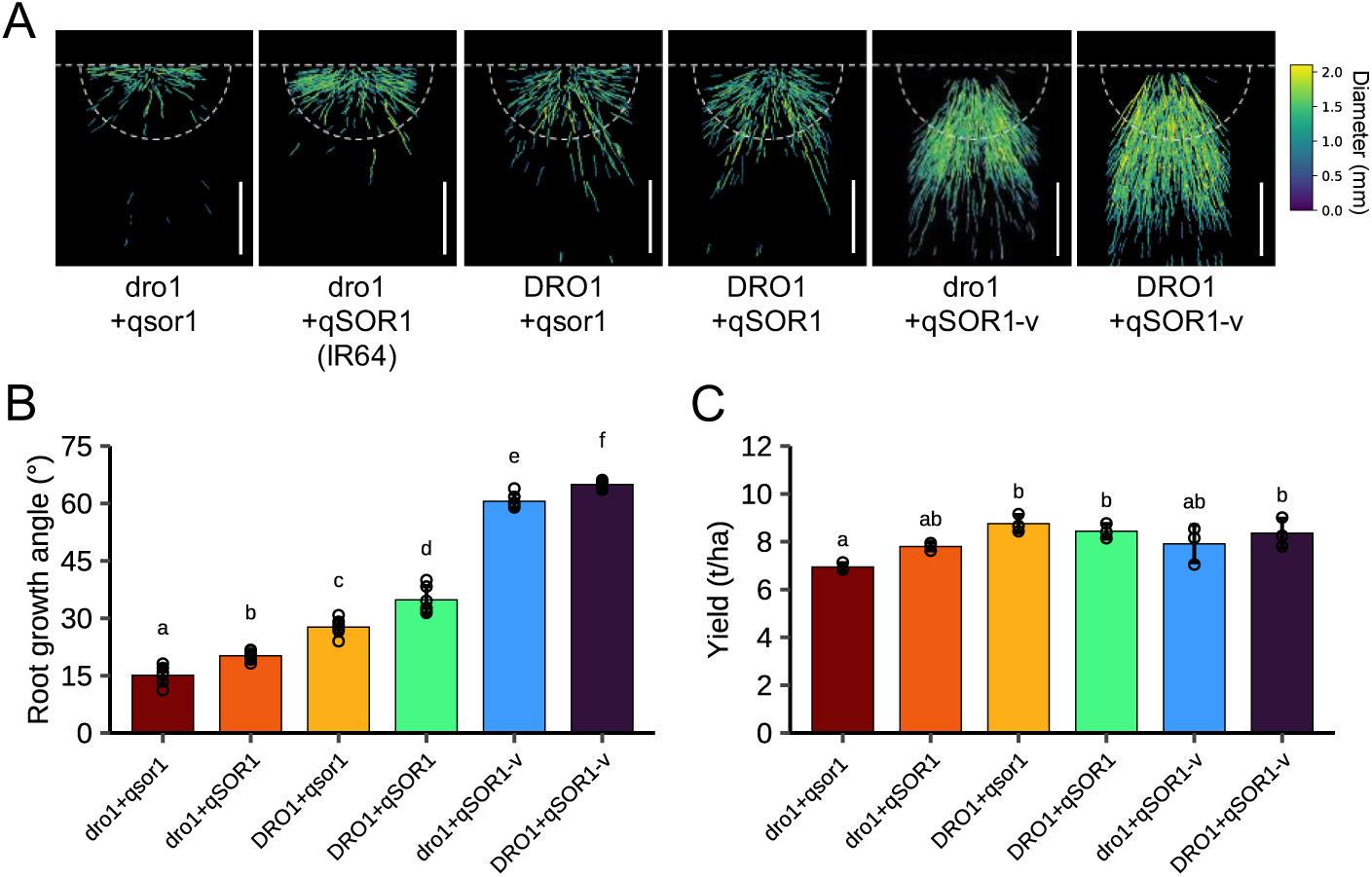
Analysis of RSA and yield in IR64 and the five ILs with different *DRO1* and *qSOR1* allele combinations grown in the paddy field. (A) Images of the RSA of rice plants at the maturity stage visualized by X-ray CT and image processing. The horizontal dashed lines and half circles indicate the soil surface and area used for the RSA measurement (radius = 8 cm), respectively. Root color represents root diameter. Scale bars, 5 cm. (B) RGA measured in (A) (*n* = 6 biological replicates). (C) Yield in each line (*n* = 3 biological replicates). Bar plots show mean values ± SD. Different lowercase letters indicate significant differences among groups (*p* < 0.05, multiple-comparison Tukey’s test).

In the IR64 background, the yield was the highest in DRO1+qsor1_PYL (+12% of IR64), and the lowest in qsor1_NIL (–11% of IR64) (Fig. 5C). Among the yield components, aboveground dry weight and harvest index were significantly positively correlated with yield (*SI Appendix*, Fig. S12F, G), with a higher harvest index in DRO1+qsor1_PYL than that in the other lines (*SI Appendix*, Fig. S11D). In the KSH background, yields did not differ significantly among the lines (*SI Appendix*, Fig. S14A). However, a significant positive correlation was observed between the RGA and yield (*SI Appendix*, Fig. S15A), indicating that vertical rooting by *qSOR1*-v contributed to a higher yield in the KSH background.

### The *qSOR1*-v with a functional *DRO1* enhances yield under drought stress

To assess the role of *qSOR1*-v in drought resistance, we studied IR64 and the five *DRO1 qSOR1* ILs under drought stress conditions in an upland field. We evaluated the RSAs of six lines grown under upland with well-watered conditions using the trench profile method (51). Vertical root distributions showed similar phenotypic patterns across the lines as observed in the paddy experiments. For example, the two lines with *qSOR1*-v showed narrower and deeper root distributions than the other lines (Fig. 6A–C).

**Figure 6.**
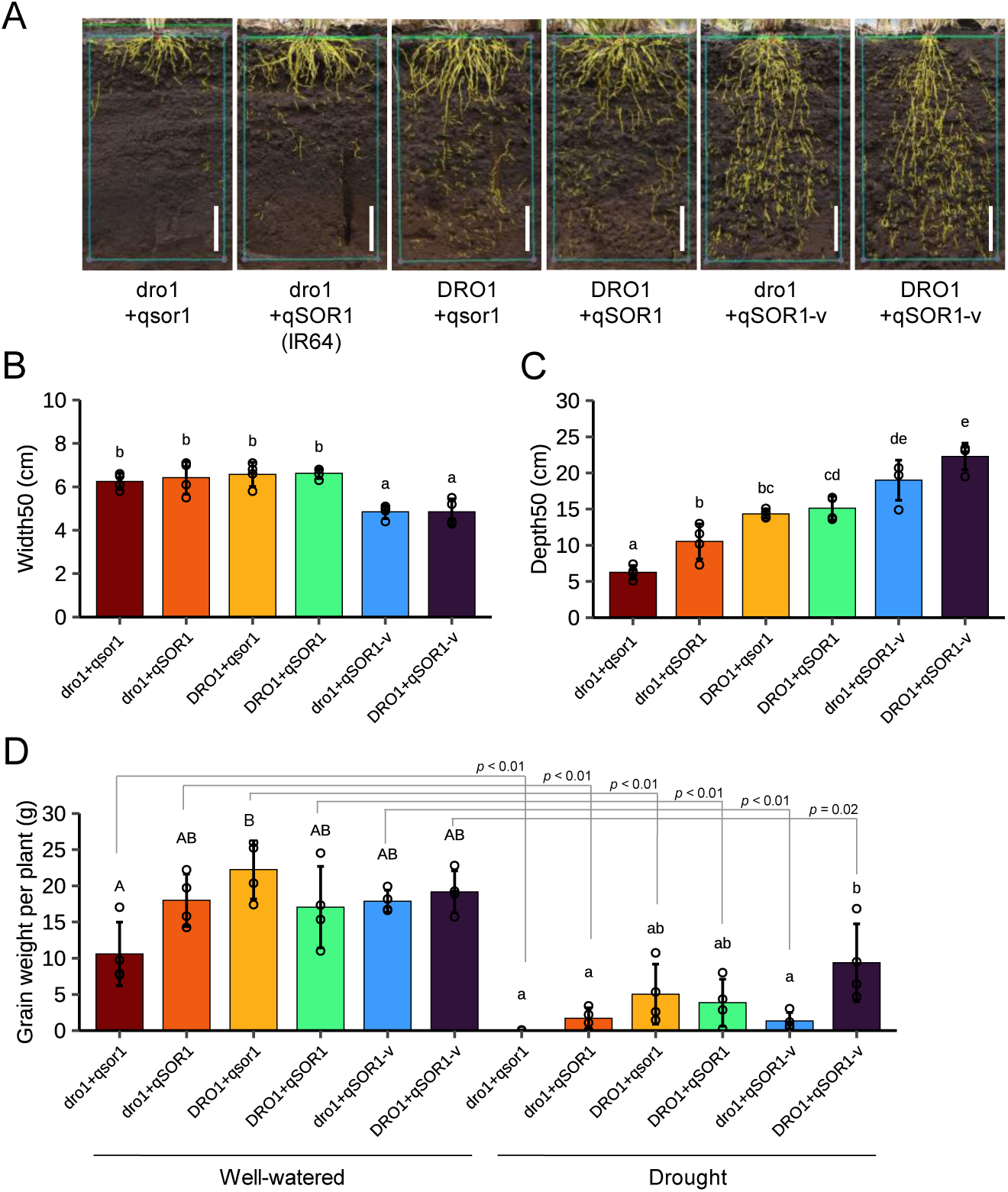
Analyses of RSA and yield in IR64 and the five ILs with different *DRO1* and *qSOR1* allele combinations grown in the upland condition. (A) Trench images of rice root systems at the maturity stage grown under well-watered conditions. The boxes indicate the area used for RSA analysis (30 cm wide and 50 cm deep). The yellow lines show roots detected by image analysis using TrenchRoot-SEG. Scale bars, 10 cm. (B, C) Horizontal (B) and vertical (C) distribution of roots measured in (A). Cumulative probabilities were calculated along both the vertical and horizontal axes. Depth50 and Width50 were defined as the depth and width at which the cumulative probability reached 0.5, respectively. (D) Grain weight under well-watered and drought conditions. Bar plots show mean values ± SD (*n* = 4 biological replicates). Different letters indicate significant differences among groups (*p* < 0.05, multiple-comparison Tukey’s test). *p* values are derived from a two-tailed Student’s *t*-test.

For the drought test, we established a greenhouse in an upland field, and drought treatment (DT) was conducted by stopping irrigation at the reproductive stage. DT lowered the soil water potential at 20 and 40 cm below the soil surface compared with well-watered conditions (WW) (*SI Appendix*, Fig. S16D). Grain weight was significantly lower in DT than in WW for all six lines (Fig. 6D). A significant positive correlation was detected between harvest index and grain weight among the six lines under DT (*SI Appendix*, Fig. S18F). Under DT, the combination of functional *DRO1* and *qSOR1*-v resulted in the highest grain weight (Fig. 6D), with a higher harvest index but no significant difference in shoot biomass between DRO1+qSOR1-v_PYL and IR64 (*SI Appendix*, Fig. S17). In contrast, ILs with non-functional *dro1* had a lower grain weight and harvest index than those with functional *DRO1*, even with *qSOR1*-v (Fig. 6D, *SI Appendix*, Figs. S17C, S18F). This suggests that vertical rooting by *qSOR1*-v with functional *DRO1* contributes to higher yields under DT.

## Discussion

By screening for *DRO1* and *qSOR1* allelic mutants, we identified *qSOR1*-v mutants with a vertical rooting phenotype (Fig. 1). Root image analyses of the ILs with *qSOR1*-v in the control chamber (Fig. 4), paddy field (Fig. 5), and upland field (Fig. 6) demonstrated that *qSOR1*-v robustly exhibited a vertical rooting phenotype across different growth conditions. As *DRO1* and *qSOR1* are involved in root gravitropism in rice (23, 42), we speculated that the vertical rooting phenotype of the *qSOR1*-v mutant was due to an altered gravity response compared to that of the WT. The root rotation assay revealed a higher accumulation of auxin with higher *OsIAA20* expression on the lower side of the root curvature in the *qSOR1*-v mutant (2792M) than in the WT (Fig. 2C, D), leading to an enhanced gravitropic response (Fig. 2A, B). Auxins play a pivotal role in the gravitropic response by regulating differential cell growth on opposite sides of the root (52). Recent studies have proposed a gravity-sensing mechanism in which LZY3 translocation from the amyloplast to the lower plasma membrane promotes the recruitment of RLD to the same region, subsequently modulating the polar localization of PIN auxin efflux carriers (37–39). The putative PEST motif, which may facilitate protein degradation through the proteasome pathway, is located in the interdomain between regions II and III (38). Deletions or amino acid substitutions in the putative PEST motif of LZY3 increased its protein abundance and promoted its accumulation on the lower side with a steeper rooting angle in *Arabidopsis* (38). In the present study, the introduction of WT *LZY3* complemented the shallow rooting phenotype of the *lzy2 lzy3* mutant (Fig. 3B, C). However, the LZY3-mCherry signal was barely detected (Fig. 3D), as previously reported (32). In contrast, the LZY3-v-mCherry signal was detected in columella cells in the root tips, even at the same expression level as in WT LZY3 (Fig. 3D, *SI Appendix*, Fig. S1D), indicating that *LZY3*-v mutation may affect protein abundance or turnover rate. Application of the protease inhibitor MG132 increased WT LZY3 fluorescence in root tips (*SI Appendix*, Fig. S7), further supporting the hypothesis that LZY3 targeted proteasome-mediated protein degradation. Further analyses are required to elucidate the function of region III of qSOR1 in rice and its orthologous proteins in relation to RLD and PINs during the root gravitropic response.

Among the five DRO1 clade subgroups identified in our phylogenetic analysis of DRO1 and LAZY1 clade proteins from the IGT family, the amino acid sequence in region III was highly conserved in the qSOR1/DRL1 and LZY2/3 subgroups of the DRO1 clade proteins (*SI Appendix*, Fig. S4, S5). Yoshihara *et al*. (33) identified five regions in the DRO1 and LAZY1 clade proteins in rice and *Arabidopsis*. Region II contains the GϕL(A/T)IGT motif, which is universally conserved in IGT family proteins (53). Region V, also known as the conserved C-terminus in LAZY1-family proteins (CCL), is well conserved among DRO1 and LAZY1 clade proteins but lacked in the TAC1 clade (32, 53). Rice LAZY1 is localized in the plasma membrane and nucleus, and deletion of its C-terminal region, including CCL, abolishes plasma membrane localization (30, 54). Rice LAZY1 interacts with OsBRXL4 through the C-terminus region of LAZY1 at the plasma membrane, which reduces nuclear localization of LAZY1 and lowers shoot gravitropism when *OsBRXL4* is overexpressed (54). *Arabidopsis* LZY3 interacts with RLD1 through the LZY3-CCL and RLD1-BRX domains, resulting in the recruitment of RLD to the plasma membrane (39). In contrast to region V (CCL), regions III and IV are less conserved among LAZY1-family proteins (*SI Appendix*, Fig. S4B, S5; ref. 33), and their functions remain largely uncharacterized (55). However, multiple alignment analysis revealed that region III, including the mutated sites of *qSOR1*-v alleles, is well-conserved among rice qSOR1, *Arabidopsis* LZY2, LZY3, and other orthologous proteins classified into the qSOR1/DRL1 and LZY2/3 subgroups (*SI Appendix*, Fig. S4B, S5). Therefore, we hypothesized that the amino acid mutation in region III of *qSOR1* orthologs expresses a similar phenotype in *Arabidopsis*. The introduction of LZY3-v with an amino acid substitution in region III (L131F), corresponding to qSOR1-v, resulted in vertical root growth in *Arabidopsis* lateral roots (Fig. 3B, C). These findings suggest that mutation of region III of the qSOR1 and LZY3 orthologs has the potential to improve RSA from shallower to deeper RSA in both monocot and dicot crops.

Drought seriously threatens crop production, particularly in rainfed regions. Deep rooting increases drought resistance in crops (13–21). The introduction of the functional allele of *DRO1* from the upland cultivar KP to IR64 resulted in deep rooting and improved yields under upland conditions (23, 24). However, the RGA in DRO1_NIL is an intermediate between IR64 and KP (23), indicating the potential for further deeper rooting. Several different RGA QTLs have been identified, including *DRO2*, *DRO3*, *DRO4*, and *DRO5*, from the crossing of KP and lowland rice cultivars (56–58), indicating that multiple genes contribute to the formation of deeper RSA in upland rice. Although deep roots are advantageous for water uptake, they incur metabolic costs for soil exploration (59). When considering the trade-off between aboveground and belowground growth, whether extreme deep rooting contributes to yield under drought conditions remains debatable. To test whether deep rooting beyond that of DRO1_NIL could lead to enhanced drought resistance, we identified the *qSOR1*-v allele that produces vertical rooting phenotypes and developed ILs with varying root distributions in their RSAs. DT significantly decreased the grain weight in IR64 and the five ILs compared to that of WW (Fig. 6D). The reduction in yield was less pronounced in DRO1_NIL than in IR64, which was consistent with the results of our previous studies (23, 24). Furthermore, the yield reduction was even less severe in DRO1+qSOR1-v_PYL, resulting in a 5.5-times higher grain weight than that in IR64 under drought conditions (*p* = 0.03, Student’s *t*-test) (Fig. 6D). In contrast, *qSOR1*-v_NIL had a significantly lower grain weight than DRO1+qSOR1-v_PYL under DT (Fig. 6D), although they showed a comparable vertical rooting phenotype (Fig. 6A–C). Because grain weight under WW was comparable in ILs possessing *qSOR1*-v with or without functional *DRO1* (Fig. 6D), unknown *DRO1* function(s) other than the RGA control, which is specifically expressed under drought conditions, may exist. These results indicate that the vertical rooting phenotype induced by the introgression of *qSOR1*-v with the combination of functional *DRO1* performed better under drought stress. Consequently, it has been proposed that steeper and deeper RSAs are more effective in mitigating yield losses associated with drought.

Although upland rice exhibits higher drought resistance than lowland rice, it generally yields less under irrigated conditions (60). This is likely due to a combination of traits in upland rice, such as a short growth duration, low photosynthesis rate, and low shoot-to-root ratio (7). As upland rice typically develops deeper and thicker root systems than lowland rice (61), the vertical rooting phenotype would negatively affect yield in irrigated paddy fields. To test this hypothesis, we evaluated the yield performance of 12 lines under a paddy condition. In this paddy field, an increased yield in DRO1_NIL (+8% of IR64, *p* = 0.04, Student’s *t*-test) was observed in the IR64 background (Fig. 5C), as previously reported (62). The yield of DRO1+qSOR1-v_PYL did not differ from that of DRO1_NIL (Fig. 5C), indicating that the vertical rooting phenotype did not confer a yield penalty under the irrigated condition. Meanwhile, DRO1+qsor1_PYL, which showed an intermediate RGA among the six lines (Fig. 5A, B), exhibited the highest yield (+12% of IR64, *p* = 0.02, Student’s *t*-test) (Fig. 5C). We observed that IR64 and the five ILs grown under WW upland conditions exhibited similar differences in yield performance compared to the results of paddy field (Fig. 5C, 6D, *SI Appendix*, Fig. S18H). These observations suggest that RSA variations play a significant role in influencing yield performance under well-watered conditions despite the differences in growth conditions between rice paddies and uplands. Notably, a significant positive correlation was detected between RGA and yield in the paddy field in the KSH background (*SI Appendix*, Fig. S15A), indicating that the ideal RSA differs by background genotype, even under the same field condition. This indicates that determining how RSA differences influence yield performance across various genetic backgrounds presents a considerable challenge for improving crop yield.

This study suggests that *qSOR1*-v is a useful allele for RSA breeding. For rice, most cultivars possess functional *DRO1* and *qSOR1* (23, 42), indicating that other genetic resources or alleles are needed to improve the root system of lowland rice to a vertical rooting phenotype similar to that of upland rice. Here, we identified *qSOR1*-v, which exhibits a robust vertical rooting phenotype across different rice varieties. No RGA-related QTLs have been detected around *qSOR1* in genome-wide association studies (50, 63, 64), and no polymorphisms have been observed in the vicinity of region III of *qSOR1* among rice genotypes in the comparative genomic database (TASUKE+; ref. 65), demonstrating the utility of *qSOR1*-v mutant allele for RSA breeding of many rice cultivars. In particular, *qSOR1*-v introgression is expected to improve drought resistance in lowland rice plants. The continuous RGA variations from shallow to vertical in the developed ILs and the parent, which were robustly expressed under different growth conditions, served as suitable materials for analyzing the RSA ideotype in different soil environments. Such information is expected to suggest the optimal RSA for the target environment, making ideotype breeding of the RSA a reality.

The conserved qSOR1 and LZY3 orthologs in angiosperms provide the potential of developing vertical rooting genotypes that enhance crop drought resistance. The increased expression levels of WT *LZY3* promoted vertical rooting; however, further overexpression resulted in more horizontal rooting than that of the WT (*SI Appendix*, Fig. S6). Notably, the overexpression of *DRO1*, *LZY3* (*AtDRO1*), and *PpeDRO1* resulted in deep rooting in rice, *Arabidopsis*, and peach, respectively (23, 35). The inducible expression of *ZmDRO1* by a synthetic ABA/drought-inducible promoter promotes deep rooting and improves grain yield under drought conditions; however, constitutively high *ZmDRO1* expression represses plant growth and grain yield (66). Although transgenes are valuable tools, their expression is often influenced by copy number, insertion site, and epigenetic regulation (67, 68). In contrast, point mutations in endogenous genes introduced through DNA marker-assisted selection or gene-editing technologies provide a more stable and heritable modification of gene functions. Consequently, *qSOR1*-v mutations in *qSOR1* and *LZY3* orthologous genes in both monocot and dicot crops have the potential to be leveraged in RSA breeding.

## Materials and Methods

### Screening and developing *qSOR1* mutant lines in rice

We screened for new mutants with non-synonymous substitutions in rice *qSOR1* (Os07g0614400) using the TILLING method (46). We used genomic DNAs from rice mutants (M1 generation, n = 3,072) obtained by treating seeds of the rice cultivar ‘Koshihikari’ (‘KSH’, *Oryza sativa* ssp. *japonica*) with *N*-methyl-*N*-nitrosourea according to a previously described method (69). We amplified *qSOR1* DNA fragments from genomic DNAs using PrimeSTAR GXL (TaKaRa, Shiga, Japan) with specific primers (#1 and #2, *SI Appendix*, Table S2). After denaturation and annealing, double-stranded DNA was cleaved by digestion with a single-strand-specific nuclease, CEL-I, at 42 °C for 60 min. Size fractionation was performed by agarose gel electrophoresis to detect mutations in *qSOR1* as cleavage products. We sequenced the PCR products from mutants in which truncated fragments were detected to determine the mutational position of the *qSOR1* gene in each mutant, using specific primers (#3 to #5, *SI Appendix*, Table S2). Sequence analysis identified four mutants containing non-synonymous substitutions in the *qSOR1* gene. The names of these four *qSOR1* mutants were 0951M, 2792M, 0909M, and 2574M. The RGA of 2574M (R204H) was identical to that of KSH. Therefore, we analyzed the other three mutants in subsequent experiments. For *DRO1*, we did not find any mutants that showed an RGA different from KSH in our population.

These three mutants contained numerous mutations throughout their chromosomes, in addition to the above-mentioned mutation in *qSOR1*. Therefore, to assess the effect of the target mutation in the *qSOR1* gene on the phenotype, we developed each backcrossed line with a homozygous allele of *qSOR1* containing the target mutation. The three mutants were backcrossed three times with KSH and then selfed. We examined the target mutation sequence using Sanger sequencing during the backcrossing and selfing processes.

### Quantification of root distribution by the basket method

To quantify root distribution in rice plants, the ratio of root number in each layer (0–10°, 10–30°, 30–50°, 50–70°, and 70–90°) was evaluated using the basket method with open stainless steel mesh baskets, as previously described (23, 24). The experiment was conducted in a glasshouse with a temperature set at 30 °C/26 °C for day/night and humidity of 50% (NICS, NARO, Tsukuba, Ibaraki, Japan). Seeds were sterilized in a fungicide (Plant Preservative Mixture; Plant Cell Technology, Washington, D.C., USA) at 15 °C for 24 h and pregerminated in water at 30 °C for two days. Germinated seeds were sown in the center of an open stainless steel mesh (7.5 cm diameter × 5.0 cm depth) filled with soil. The basket was placed on a polyvinyl chloride (PVC) pipe in a container filled with 1/2 Kimura B nutrient solution (70). At 35 DAS, the number of roots that had penetrated the mesh in each layer was determined and divided by the total number of roots that had penetrated the mesh. The ratio of deep rooting was defined as the number of roots in the 50– 90° layer divided by the total number of roots.

### Evaluation of root gravitropic curvature

The gravitropic curvature of the seminal root was evaluated in rice seedlings after rotating the plants, as previously described (23, 42). Seedlings were grown on 0.4% agarose in the dark at 28 °C for 2 d. Subsequently, plants were rotated from the normal vertical axis to the horizontal axis for 4 h, and images were taken with a digital camera (D5600; NIKON Co., Tokyo, Japan). The root angle of curvature was measured using the ImageJ software (https://imagej.net/ij/).

### Measurement of the lateral root tip angle in *Arabidopsis*

We used *Arabidopsis thaliana* accession Columbia-0 (Col-0) as WT *Arabidopsis*. The T-DNA-inserted loss-of-function mutant lines of *LZYs* used in this study, *lzy3*, *lzy2 lzy3*, and *lzy2 lzy3 lzy4*, were generated previously (32, 71). The seeds were soaked in 70% EtOH for 5 min and sterilized by inversion in 5% hypochlorous acid and 1% SDS for 5 min. The sterilized seeds were then pregerminated in water at 4 °C for 2 d. Germinated seeds were sown on a 1% agarose containing 1/2 MS medium. Seedlings were grown under a long-day condition (16L/8D) at 22 °C in a growth chamber. After 12 d of sowing, root images were captured using a digital camera (D5600; NIKON Co., Tokyo, Japan). The angles between the lateral root tip and gravity vector were measured using ImageJ (https://imagej.net/ij/).

### RNA isolation and expression analysis by qRT-PCR

Total RNA was isolated from the roots of rice and *Arabidopsis* plants grown for 2 and 12 d, respectively, as described above, using the RNeasy Plant Mini Kit (Qiagen, Hilden, Germany) or TRIzol Reagent (Invitrogen, Carlsbad, CA, USA) according to the manufacturer’s instructions. First-strand cDNA was synthesized using SuperScript II reverse transcriptase (Invitrogen, Carlsbad, CA, USA). qRT-PCR was performed using the THUNDERBIRD SYBR qPCR Mix (TOYOBO, Osaka, Japan) and ViiA7 Real-Time PCR System (Life Technologies, Carlsbad, CA, USA). Target gene expression was normalized to that of *OsUBQ10* (Os02g0161900) and *ACT2* (At3g18780). All primers used for qRT-PCR are listed in *SI Appendix*, Table S2 (#6 to #15).

### Quantification of endogenous IAA amount during root gravitropic curvature

To quantify the amount of endogenous IAA in the seminal root during root gravitropic curvature, rice seedlings were grown and rotated as described above. After 1 h of rotation, the upper and lower sides of root segments at 0.5–2.5 mm from the root tip were sampled by splitting the roots with a needle under a microscope (STZ-161-TLED-1080M; Shimadzu Corporation, Kyoto, Japan). The sampled root segments were used to quantify endogenous IAA and *OsIAA20* expression. The collected root segments were frozen in liquid nitrogen immediately after harvest and stored at –80 °C until use. After lyophilization, the dry weights of the samples were measured. Then, the dried samples were pulverized under ice-cold conditions by adding 600 µL of 80% methanol containing 0.1% formic acid per mg of samples using a Multi-Beads Shocker MB1050 (Yasui Kikai, Osaka, Japan) at 2,500 rpm for 40 s. The extract was centrifuged at 5,000 × *g* for 5 min at 4 °C and filtered through a 0.2-µm pore size filter. Following centrifugal concentration of a 450 µL aliquot of the filtrate, the residue was dissolved in 15 µL of 30% acetonitrile containing 10 pg/µL IAA-d_5_ under ice-cold conditions. After centrifugation at 12,000 × *g* for 10 min at 4 °C, the supernatant was analyzed by liquid chromatography/mass spectrometry (LC/MS) using a Vanquish Flex ultra-high-performance liquid chromatography system coupled with an Orbitrap Q Exactive Focus instrument (Thermo Fisher Scientific, MA, USA). Five microliters of the supernatant were injected and separated on an ODS column (Ascentis Express 90 Å AQ-C18 50×2.1 mm I.D., 2.7 µm; Merck, Darmstadt, Germany) at a flow rate of 0.4 mL/min. The solvent program consisted of 10% (0–1 min), 10–100% (1–5 min), and 100% (5–7.5 min) acetonitrile containing 0.1% formic acid in H_2_O in linear gradient mode. The temperatures of the preheater and column oven were set at 50 °C and maintained in the forced mode. The operating parameters of the mass spectrometer are presented in the *SI Appendix* (Table S3). The acquired data were analyzed using FreeStyle (version 1.7.85.0) and QuanBrowser Xcalibur ver 4.3.73.11 (Thermo Fisher Scientific, Waltham, MA, USA).

### Plasmid construction and plant transformation in *Arabidopsis*

*LZY3p*-*LZY3*-*mCherry*/pZErO and *LZY3p*-*LZY3*-*mCherry*/pGWB501 vectors were developed based on previous studies (32, 38). To obtain transformants of *LZY3*-v (L131F) in the *lzy2 lzy3* background, point mutations were introduced by PCR amplification with *LZY3p*-*LZY3*-*mCherry*/pZErO as a template using specific primers (#16 and #17, *SI Appendix*, Table S2), followed by self-ligation. Mutated *LZY3*-v was introduced into the AvrII/PacI sites of the *LZY3p*-*LZY3*-*mCherry*/pGWB501 construct. The resulting plasmid vector was introduced into *Agrobacterium tumefaciens* strain GV3101 by electroporation. *LZY3* or *LZY3*-v was introduced into *lzy2 lzy3* using the *Agrobacterium*-mediated floral dip method (72). Homozygous transgenic lines for each transgene were established at T3. Individuals carrying single-copy *LZY3* or *LZY3*-v transgenes were selected from the segregating progeny derived from the seeds of a heterozygous parent by expression analysis of the transgenes.

### Microscopic analysis of reporter gene expression in *Arabidopsis* root tips

*Arabidopsis* plants, *LZY3p*-*LZY3*(-v)-*mCherry*/*lzy2 lzy3*, were grown on 1% agarose containing the 1/2 MS medium as described above. After 12 d of sowing, the main root tips were sampled and observed under a laser scanning microscope (LSM710; Carl Zeiss, Oberkochen, Germany). mCherry fluorescence was excited at 561 nm and detected at 570–620 nm. For the chemical treatment, the intact roots were incubated in the 1/2 MS medium containing 50 μM MG132 for 16 h.

### Subcellular localization analysis in rice protoplast

The ORF sequence of *qSOR1* was subcloned into pENTR4 (Invitrogen, Carlsbad, CA, USA), as previously described (42). The point mutation in *qSOR1* (L141F) was introduced by PCR amplification with *qSOR1*/pENTR4 as a template using specific primers (#18 and #19, *SI Appendix*, Table S2), followed by self-ligation. The mutated *qSOR1*-v was then transferred to pSAT6-DEST-EGFP-C1(73) using the Gateway LR reaction (Invitrogen, Carlsbad, CA, USA). Next, 2×*35Sp-qSOR1*(-v)-*EGFP* and 2×*35Sp-EGFP* were transfected into the protoplasts of young rice seedlings, as previously described (42). Briefly, the 100 μL protoplast suspension (total 200,000 cells) isolated from 1-week-old seedlings was transfected with the 10 μL plasmid vector (total 10 μg). After an overnight incubation, EGFP fluorescence was observed using a laser scanning microscope (LSM710; Carl Zeiss, Oberkochen, Germany). EGFP fluorescence was excited at 488 nm and detected at 510–530 nm.

### Development of near-isogenic lines (NILs) and pyramiding lines (PYLs) with multiple allele combinations of *DRO1* and *qSOR1*

To investigate the influence of *DRO1* and *qSOR1* combinations on RSA and agronomic traits, we developed five rice lines with two genetic backgrounds, KSH and IR64 (*O. sativa* ssp. *indica*). These lines incorporated three alleles of *qSOR1* (non-functional, functional, and enhanced functional) and two alleles of *DRO1* (non-functional and functional) (*SI Appendix*, Table S1).

To establish the five NILs and PYLs in the genetic background of KSH, we used the 2792M line as the NIL for *qSOR1*-v (qSOR1-v_NIL) and the 0909M line as the NIL for *qsor1* (qsor1-NIL). We also used one of the IR64 chromosome segment substitution lines (CSSLs) in the KSH background (74), in which the vicinity of the *DRO1* locus was replaced with an IR64 allele (*dro1*), as a donor to generate dro1_NIL. Using SSR markers, we selected a homozygous line from the F_2_ progeny of a self-pollinated plant between CSSL and KSH, in which only in the vicinity of the *DRO1* locus was the IR64 allele using SSR markers. To generate dro1+qsor1_PYL and dro1+qSOR1-v_PYL, we selected the desired genotype combinations of the *DRO1* and *qSOR1* loci from the F_3_ progeny of plants self-pollinated between dro1_NIL and qsor1_NIL and between dro1_NIL and qSOR1-v_NIL, respectively. We examined the target mutation sequence using Sanger sequencing during the backcrossing and selfing processes.

To establish the five NILs and PYLs in the genetic background of IR64, we used the three IR64 lines reported by Kitomi *et al*. (42). In the present study, we developed two new lines containing the *qSOR1*-v alleles. We backcrossed qSOR1-v_NIL in a KSH background with IR64 four times, followed by repeated selfing, and selected a line with the homozygous allele of *qSOR1*-v (qSOR1-v_NIL) from the F_5_ progeny. We crossed a BC_4_F_1_ plant obtained during the selection process of qSOR1-v_NIL to the DRO1-NIL, followed by repeated selfing, and selected a line with homozygous alleles of *DRO1* and *qSOR1-v* (DRO1+qSOR1-v_PYL) from the F_5_ progeny. We used SNP markers to confirm the genetic background and SSR markers to narrow down the target regions during the selection process for both lines. All DNA marker-assisted selection using SSR and SNP markers was conducted according to previously described methods (58).

### Time-series 3D root imaging using X-ray CT

We evaluated the temporal changes in 3D RSA development in 12 rice lines (KSH, five KSH ILs, IR64, and five IR64 ILs described in *SI Appendix*, Table S1) through 3D image analysis using X-ray CT. Rice was grown in pots under controlled artificial conditions, as previously described (49, 75). The environmental conditions in the growth chamber (Nippon Medical & Chemical Instruments Co., Osaka, Japan) were 14 h of light and 10 h of dark (NICS, NARO, Tsukuba, Ibaraki, Japan). The temperature was 30 °C/25 °C, and the humidity was 50%/60% for day/night, respectively. We used an automated irrigation system that allowed water to be supplied from the bottom of the pot (iPOTs; ref. 49). One iPOT unit can manage six pots simultaneously. Six plants were arranged in each line in a randomized complete block design (RCBD). We controlled 8 cm of water from the bottom of the pot during the cultivation period using iPOTs. Each pot was filled with a soil-like substrate (Profile Greens Grade; PROFILE Products, Buffalo, IL, USA). Before sowing, the soil-filled pot was soaked in a modified Kimura B nutrient solution (70), as previously described (49). After sterilizing and germinating the seeds, we sowed three germinated seeds from each line in the center of each pot. Approximately one week after sowing, we selected the average established seedlings and removed the other two.

We conducted CT scanning of each plant four times from the second to the fifth week post-sowing using an X-ray CT system (inspeXio SMX-225CT FPD HR; Shimadzu Corporation, Kyoto, Japan), as previously described (48). The CT scanning conditions were as follows: tube voltage 225 kV, tube current 500 μA, 1.0-mm-thick copper filter, and 5 min of scanning time. A total of 860 tomography images were obtained, each with a 1024 × 1024 pixel resolution and 16-bit depth. We visualized the roots from the CT image volumes using RSAvis3D (48) and vectorized them using RSAtrace3D (76). We obtained three root parameters (RDI, average RGA, RN, and TRL) using the RSAtrace3D (76).

### Measurement of yield and yield components in a paddy field

We investigated the yield and related components of 12 rice lines (*SI Appendix*, Table S1) in paddy fields according to previously described methods (62). As a basal dressing for the paddy, we applied PK compound fertilizer and controlled-release fertilizers at N:P_2_O_5_:K_2_O = 12:12:9 kg 10a^−1^. The three controlled-release N fertilizers (LP40, LPs100, and LP140) released 80% of the total nitrogen at uniform rates for up to 40, 100, and 140 d, respectively. We seeded the nursery box on April 15, 2024, followed by transplanting established seedlings with one plant per hill in a paddy field at the Kannondai site, Tsukuba, Ibaraki, Japan (36°01′ N, 140°06′ E) on May 15, 2024. With a spacing of 30 cm between rows and 18 cm between hills, we planted 12 plants per row in each line for a total of 72 plants per plot. Within each KSH and IR64 genetic background replication, we arranged six lines in the RCBD with three replicates.

The heading date for each line was defined as the date at which 50% of the plants were heading. We harvested 24 plants from each line in bulk, followed by threshing and selection using a fan. The samples were then dried in an oven at 80 °C for over 3 d to calculate yield. Before harvesting, we measured the culm length, panicle length, and panicle number of six plants from each line. To measure the yield components, we selected two plants with an average number of panicles from their respective lines. After harvesting, yield components were measured as described by Arai-Sanoh *et al*. (62).

### Evaluation of 3D root system architecture in paddy field

To observe the 3D RSA at the mature stage of the 12 rice lines (*SI Appendix*, Table S1) in paddy fields, we conducted CT scanning and imaging of soil monolith blocks collected from paddy fields, following previously reported methods (50). After harvesting, two hills representing the average number of tillers were selected from each plot for each line. As previously described, monolith blocks were sampled from paddy fields (50). We used a pile driver to drive a custom-made steel cylinder monolith (diameter of 16 cm and length of 40 cm) approximately 25 cm below the soil surface. The monolith was pulled out, and a 20 cm-long monolith block was collected.

The monolithic blocks were scanned using X-ray CT. The CT scanning conditions were as follows: tube voltage 225 kV, tube current 500 μA, and 5 min of scanning time. The root segments were visualized using RSAvis3D (48) and RSApaddy3D (50). Root parameters, including the average RGA, average RD, and RN, were calculated and exported from RSApaddy3D.

### Drought stress test in an upland field

To investigate the effects of differences in RSA on drought resistance, we used IR64 and five IR64 ILs. We established a greenhouse (7.2 m wide × 30 m long) in an upland field to establish well-watered (WW) and drought stress conditions. As a basal dressing for the uplands, we applied NPK compound fertilizers at N:P_2_O_5_:K_2_O = 5.6:12.8:5.6 kg 10a^−1^. Topdressing (N and K, 1.0 kg 10a^−1^) was performed eight weeks after sowing (WAS). With a spacing of 30 cm between the rows and 15 cm between the hills, we planted 15 plants per row in each line, totaling 45 plants per plot. Within each replicate, we arranged six lines in the RCBD with four replicates. Rice seeds were sown three times per hill on May 27, 2024. Approximately 4 WAS, we selected the average seedlings and removed two seedlings.

The drought treatment (DT) was applied from panicle initiation (12 WAS) until the heading date (16 WAS) with reference to IR64, during which irrigation was suspended in the DT plots. From 8 WAS, we periodically monitored for panicle initiation in IR64 grown in border plots. In contrast, the WW plots were irrigated biweekly until harvest. At 22 WAS, we harvested the bulk of 10 plants from each line, followed by threshing and fan selection. The samples were then dried in an oven at 80 °C for over 3 d to calculate grain weight.

We monitored the soil water potential at depths of 20 and 40 cm as well as air conditions (temperature and humidity) in the WW and DT plots using soil water potential sensors and temperature and humidity sensors (TEROS-21 and ATMOS14; METER Group, WA, USA), respectively. Environmental data were stored in a data logger (Em50; METER Group, WA, USA).

### Evaluation of vertical root distribution in an upland field using trench profile method

To investigate the differences in vertical root distribution among the six lines of IR64, we conducted a trench profile method and 2D image analysis according to previously described methods (51). Because it was difficult to manually excavate and evaluate root distribution in the greenhouse, rice plants were grown in the upland field using the same experimental design as in the greenhouse. At harvest, we observed the root distribution using the trench profile method. We used a backhoe to cut down to a few centimeters in front of each hill, excavating at least 150 cm wide and 100 cm deep. Approximately 1–2 cm of the soil layer on the profile wall was flushed with water using an agricultural nebulizer. The exposed roots on the profile walls of three plants in each line were imaged using digital cameras (D5600; NIKON Co., Tokyo, Japan). We calculated the vertical and horizontal centroids of root distribution (Depth50 and Width50) from profile images using TrenchRoot-SEG (51). Environmental data for the upland field were obtained in a manner similar to that for the greenhouse.

## Acknowledgments

We thank M. Morita (NIBB) for kindly providing the *LZY3p-LZY3-mCherry*/pZErO and *LZY3p-LZY3-mCherry*/pGWB501 vectors and *lzy3*, *lzy2 lzy3*, and *lzy2 lzy3 lzy4* mutants; H. Imaizumi (NIAS, NARO) and N. Tanaka (NICS, NARO) for their technical support with the microscopy experiments; N. Inagaki, W. Tsuchiya, and N. Suzuki (Advanced Analysis Center, NARO) for their valuable advice in the protein experiment; H. Sakakibara (Nagoya U.) and M. Kojima (RIKEN) for their valuable advice in auxin quantification; S. Sakamoto (AIST) for his valuable advice on the protoplast experiment; M. Yamazaki for conducting the mutant screening; and Y. Fukuda, H. Tanaka, K. Onoe, N. Kanno, C. Kawashima, and the technical support section of NARO for their technical assistance in the plant phenotyping and field trials. This work was supported by JST CREST (JPMJCR17O1), Cabinet Office, Government of Japan; the Moonshot Research and Development Program for Agriculture, Forestry, and Fisheries (funding agency: Bio-oriented Technology Research Advancement Institution, JPJ009237), and JSPS KAKENHI (19H02936).

