## Supporting_information for "Vertical rooting caused by enhanced functional allele of *qSOR1* improves rice yield under drought stress"

**This PDF file includes:**

Figures S1 to S18
Tables S1 to S3
SI References

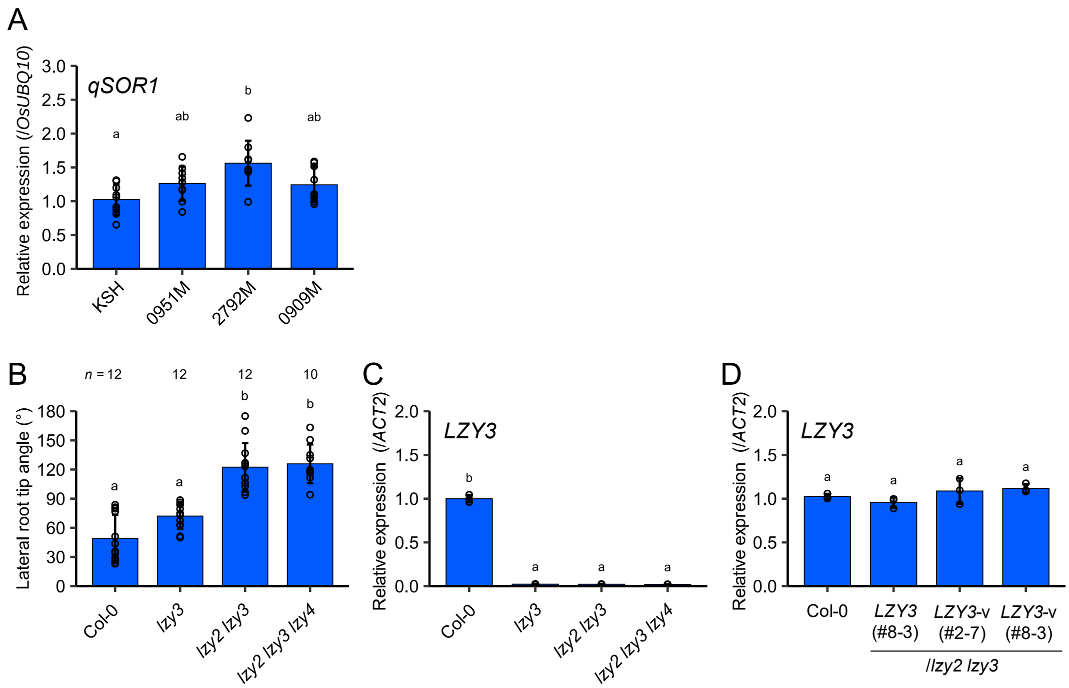


**Figure S1.** Analysis of *qSOR1* and *LZY* mutants/transgenic lines. (A) *qSOR1* expression in seminal root tips of KSH and the mutants ($n$ = 9 biological replicates). The expression of *qSOR1* was normalized to that of *OsUBQ10*. (B) Lateral root tip angle in Col-0 and the *lzy* mutants. The angle of each root tip relative to the gravity vector was measured. (C, D) *LZY3* expression in whole roots of (C) Col-0 and the *lzy* mutants and (D) *lzy2 lzy3* transgenic lines harboring either wild-type *LZY3* or v-type mutated *LZY3* (*LZY3*-v, L131F) ($n$ = 3 biological replicates). The expression of *LZY3* was normalized to that of *ACT2*. Bar plots show mean values ± SD. Different lowercase letters indicate significant differences among groups ($p < 0.05$, multiple-comparison Tukey's test).

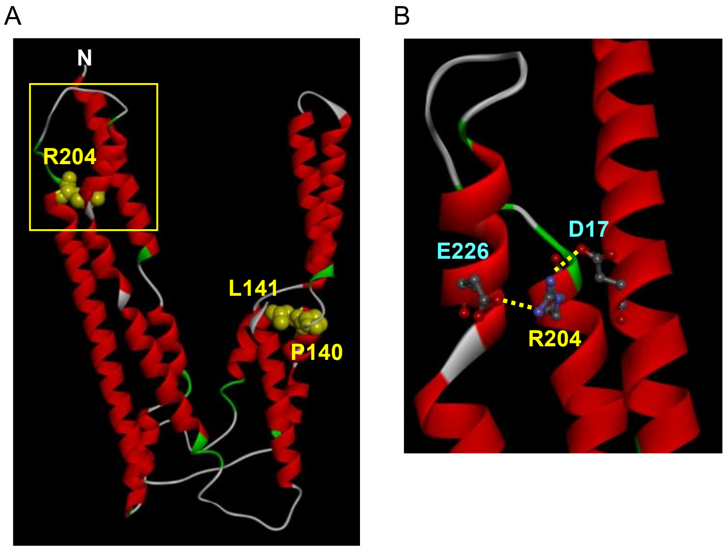


**Figure S2.** (A) 3D model of the qSOR1 protein in KSH generated using I-TASSER (1–3). The positions of amino acid residues mutated in the mutants are indicated in yellow. (B) Enlarged view of the boxed area in (A). The positions of the amino acid residues that interact with R204 via salt bridges are indicated in light blue.

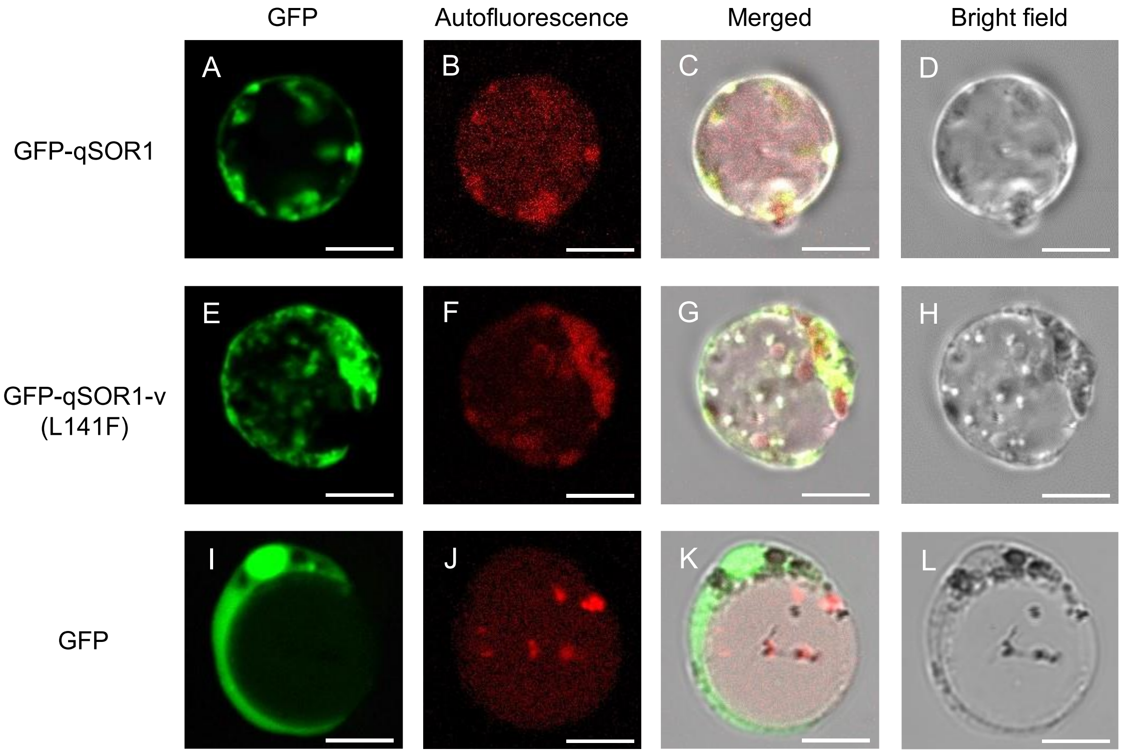


**Figure S3.** Subcellular localization of qSOR1 in rice protoplasts transformed with (A–D) 2×*35Sp-qSOR1-EGFP*, (E–H) 2×*35Sp-qSOR1*-v(L141F)-EGFP, and (I–L) 2×*35Sp-EGFP*. Panels show (A, E, I) EGFP fluorescence, (B, F, J) plastid autofluorescence, (C, G, K) merged images, and (D, H, L) bright field images. Scale bars, 10 μm.

A

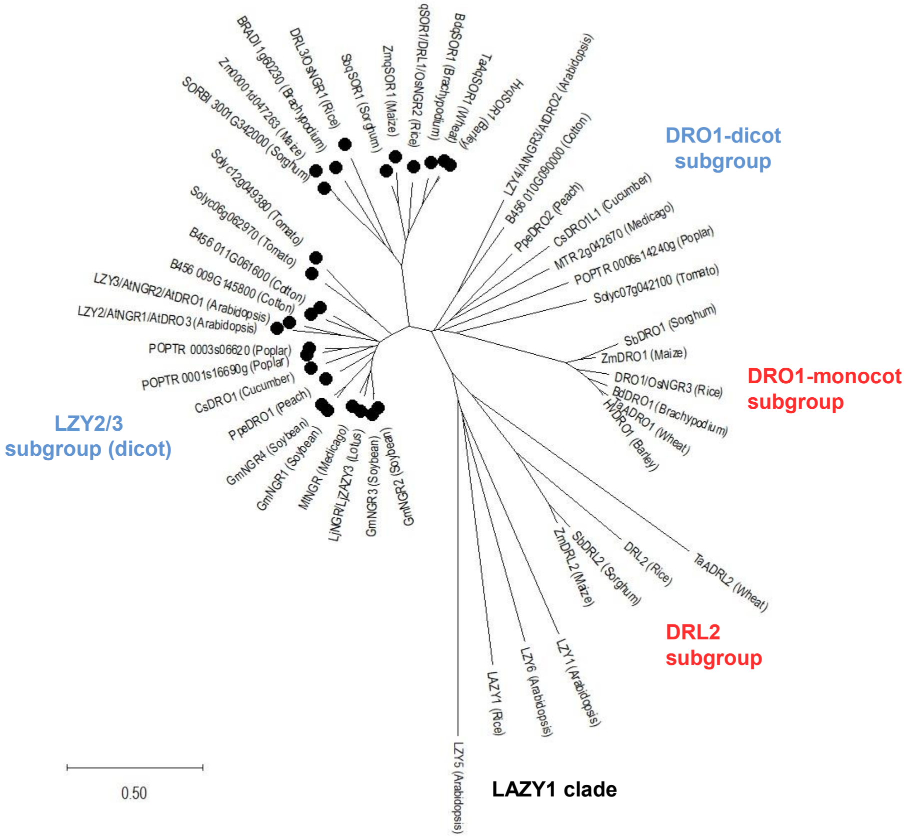


B

| | Region III | |
|---|---|---|
| qSOR1/DRL1/OsNGR2 (Rice) | L L L P L D R F L - - - - - N C P - - - - S S L E V D R R V | 157 |
| DRL3/OsNGR1 (Rice) | S G L P L D M F L - - - - - N C P - - - - S S L E V D R R A | 244 |
| LZY2/AtNGR1/AtDRO3 (Arabidopsis) | K N L P L D R F L - - - - - N C P - - - - S S L E V D R R I | 145 |
| LZY3/AtNGR2/AtDRO1 (Arabidopsis) | A N L P L D R F L - - - - - N C P - - - - S S L E V D R R I | 147 |
| LAZY1 (Rice) | V D C P L Q G F L F G S P V S D V E S R P E Y L Q A P R D S | 221 |
| DRO1/OsNGR3 (Rice) | - - - - - - - - - - - - - - - - - - - - - - - - - - - - - - | 110 |
| DRL2 (Rice) | - - - - - - - - - - - - - - - - - - - - - - - - - - - E D D | 100 |
| LZY1 (Arabidopsis) | M K F P L Q E Y F F G S L I E L P - - - - E S K I A G K K D | 207 |
| LZY4/AtNGR3/AtDRO2 (Arabidopsis) | - - - - - - - - - - - - - - - - - - - - - - - - - - - - - - | 61 |
| LZY5 (Arabidopsis) | K A C F S S T F T L R R L N S F R R G - - H L L R S M A K I | 93 |
| LZY6 (Arabidopsis) | N E C P L Q D Y D - - - - - L F R - - - - S S I E L - - - - | 147 |

**Figure S4.** Phylogenetic analysis of qSOR1 and its orthologous proteins. (A) Phylogenetic tree constructed from full-length protein sequences of DRO1 and LAZY1 clade proteins from both monocots and dicots using MEGA11. The proteins marked with a filled circle share the conserved sequences in region III. The scale bar indicates evolutionary distance based on amino acid substitutions. (B) Multiple sequence alignment of the vicinity of region III from qSOR1 orthologs in rice and *Arabidopsis*, belonging to DRO1 and LAZY1 clades. Amino acid residues highlighted with a gray background are conserved. The two amino acid residues shown in red represent mutation sites in the two *qSOR1*-v mutants: P140S in 0951M and L141F in 2792M.

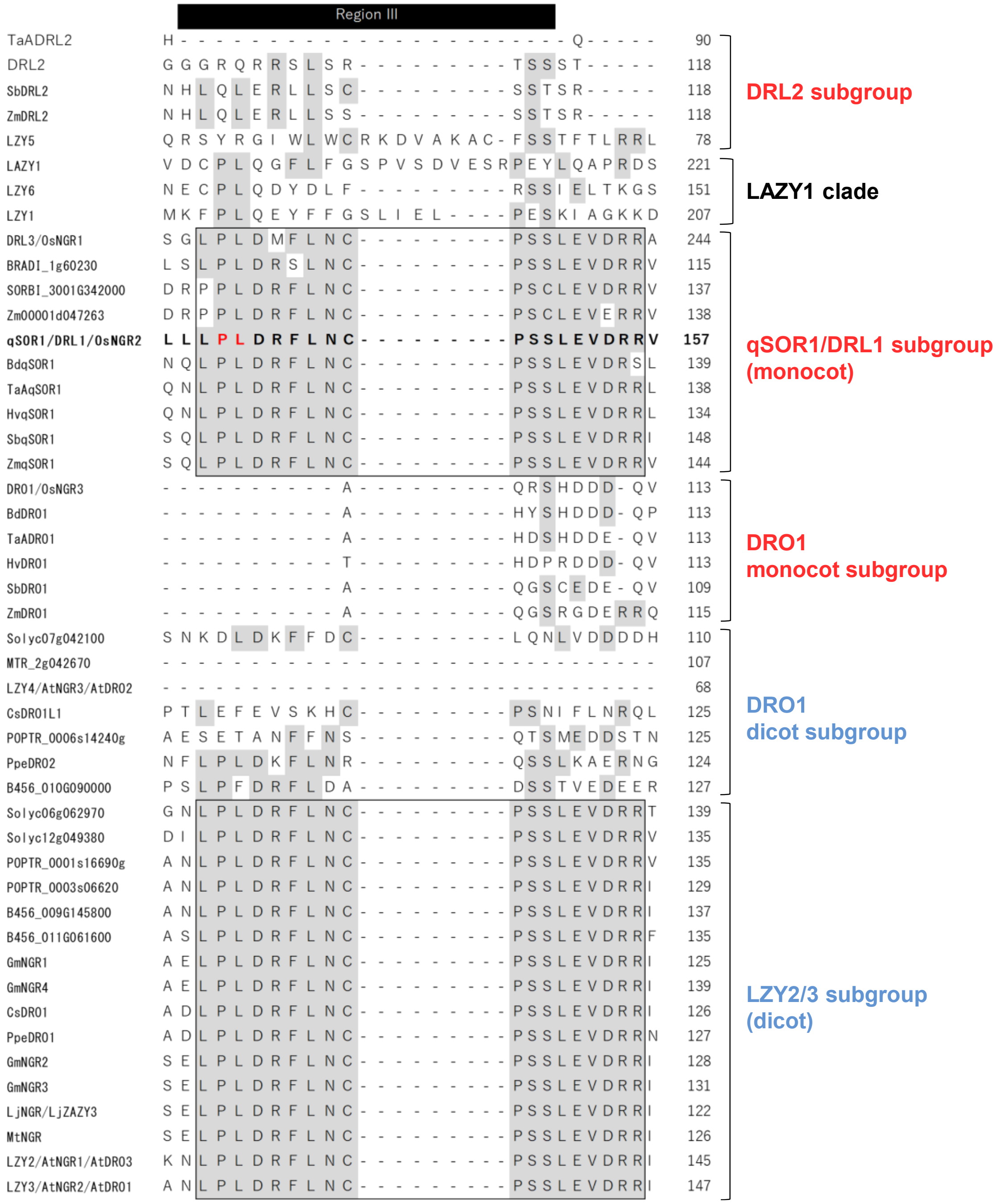


**Figure S5.** Multiple sequence alignment of the vicinity of region III from qSOR1 orthologous proteins in monocots and dicots belonging to DRO1 and LAZY1 clades. Amino acid residues highlighted with a gray background are conserved. The two amino acid residues shown in red represent mutation sites in the two *qSOR1*-v mutants (P140S in 0951M and L141F in 2792M). The black boxes indicate the conserved regions shared by qSOR1/DRL1 and LZY2/3 subgroups.

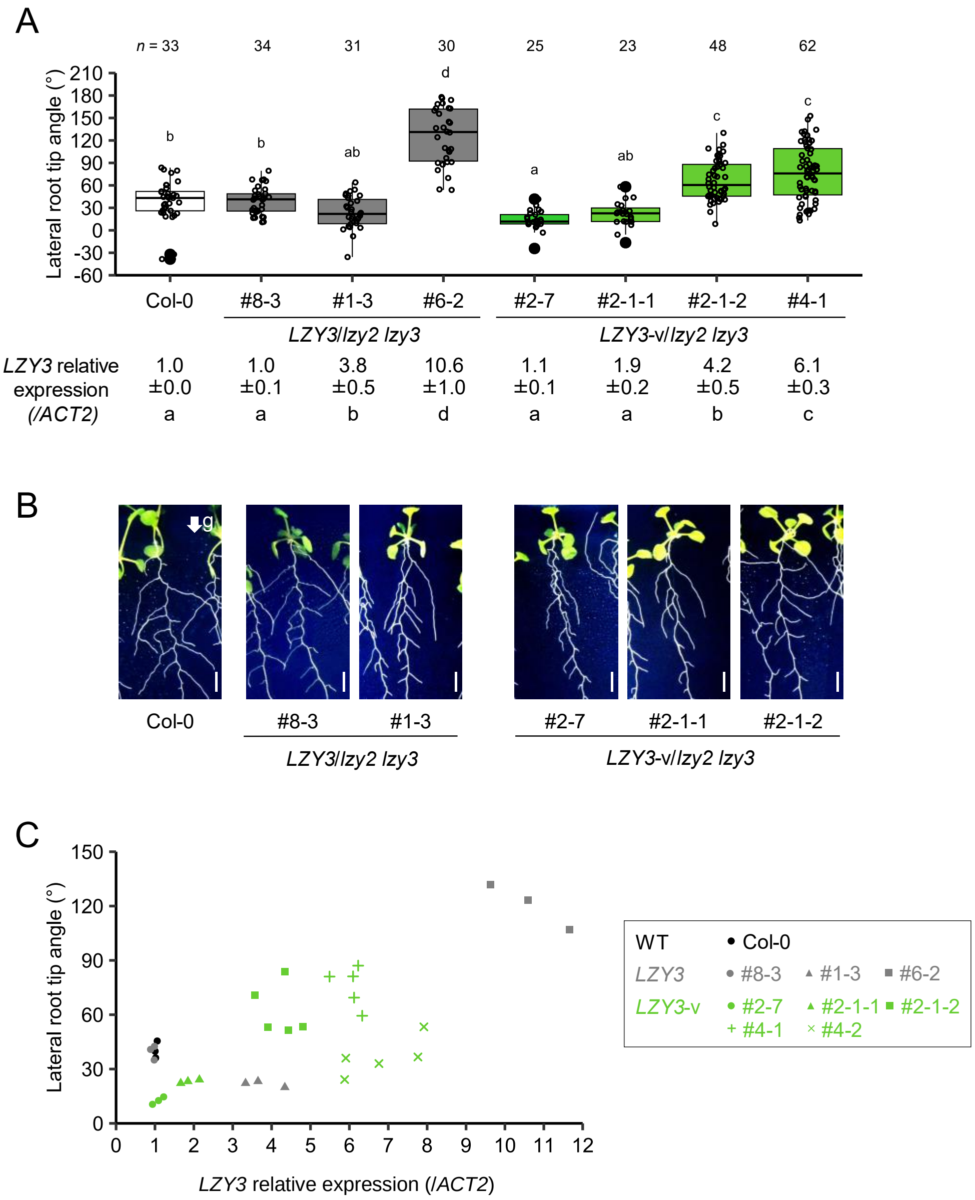


**Figure S6.** Relationship between *LZY3* expression levels and root growth angle in *Arabidopsis*. (A) Lateral root tip angles in Col-0 and *lzy2 lzy3* transgenic lines harboring either wild-type or v-type mutated *LZY3* (*LZY3*-v, L131F). Relative expression levels of *LZY3* in each line are shown below the graph (*n* = 3–5 biological replicates). The expression of *LZY3* was normalized to that of *ACT2*. Boxes represent the first quartile, median, and third quartile. Whiskers show the range of nonoutlier values. Filled circles indicate outlier values less than the first quartile or greater than the third quartile by 1.5 times the interquartile range. Different lowercase letters indicate significant differences among groups ($p < 0.05$, multiple-comparison Tukey's test). (B) Images of the representative lines in (A). The arrow indicates the direction of the gravitational force. Scale bars, 5 mm. (C) Scatter plot showing the relationship between lateral root tip angle and *LZY3* relative expression. Each dot represents an individual plant.

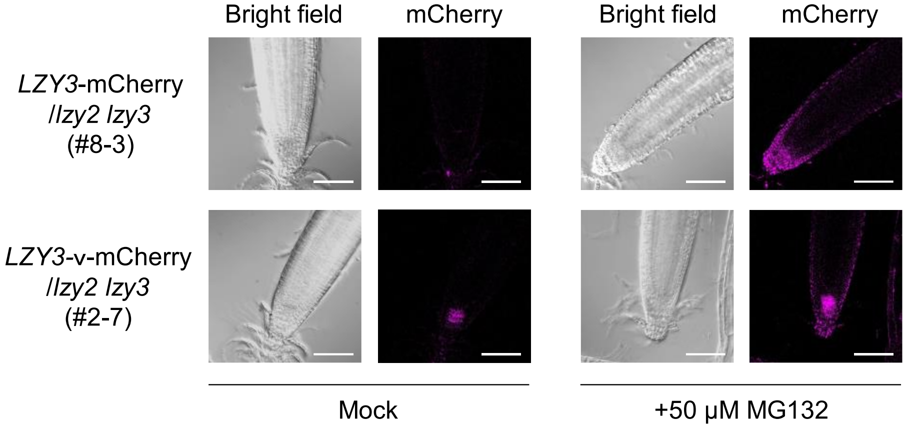


**Figure S7.** Expressions of wild-type *LZY3* and *LZY3*-v (L131F) fused to the mCherry fluorescent protein in main root tips of *lzy2 lzy3* mutant, with or without the treatment of the proteasome inhibitor, MG132. Root parts of each line were treated with 50 µM MG132 for 16 h prior to the microscope observation. Scale bars, 50 µm.

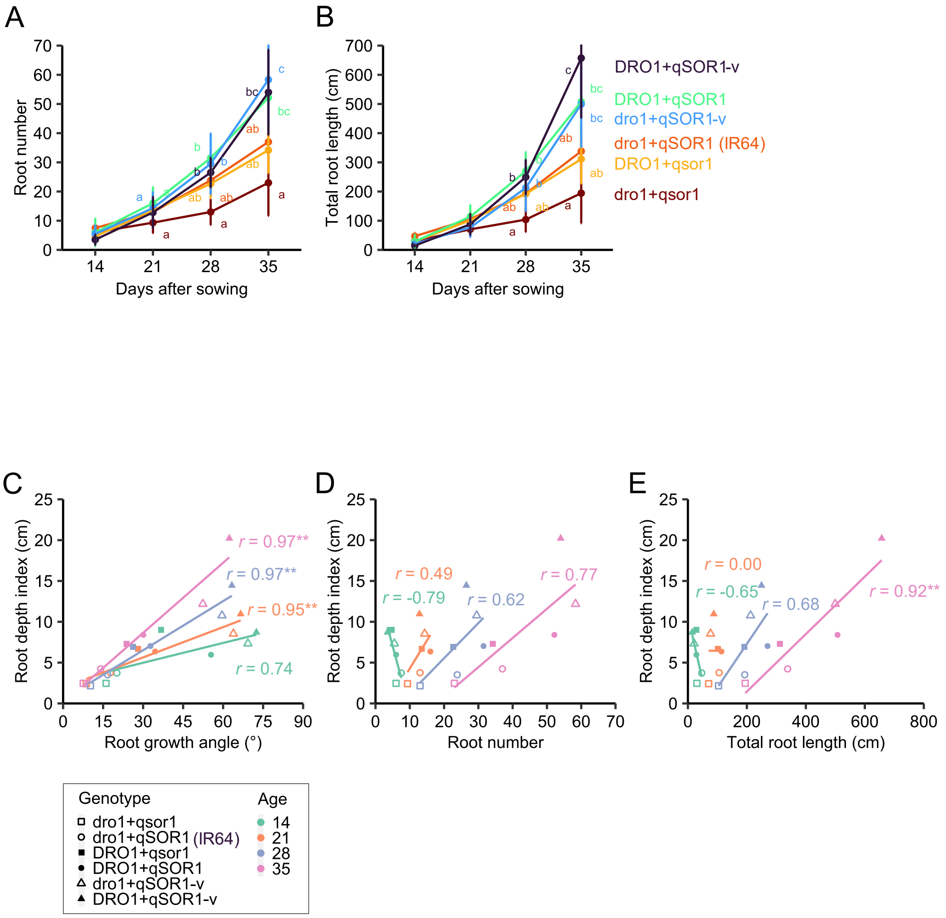


**Figure S8.** RSA analysis of IR64 and the five ILs with different *DRO1* and *qSOR1* allele combinations grown in a plant pot under a controlled chamber. (A, B) Time-course analyses of (A) root number and (B) total root length. Line charts show mean values ± SD ($n$ = 6 biological replicates). Different lowercase letters indicate significant differences among groups ($p < 0.05$, multiple-comparison Tukey's test). (C–E) Relationships between root depth index and (C) root growth angle, (D) root number, and (E) total root length across the six lines at different plant ages. $r$, Pearson's correlation coefficient (**$p < 0.01$).

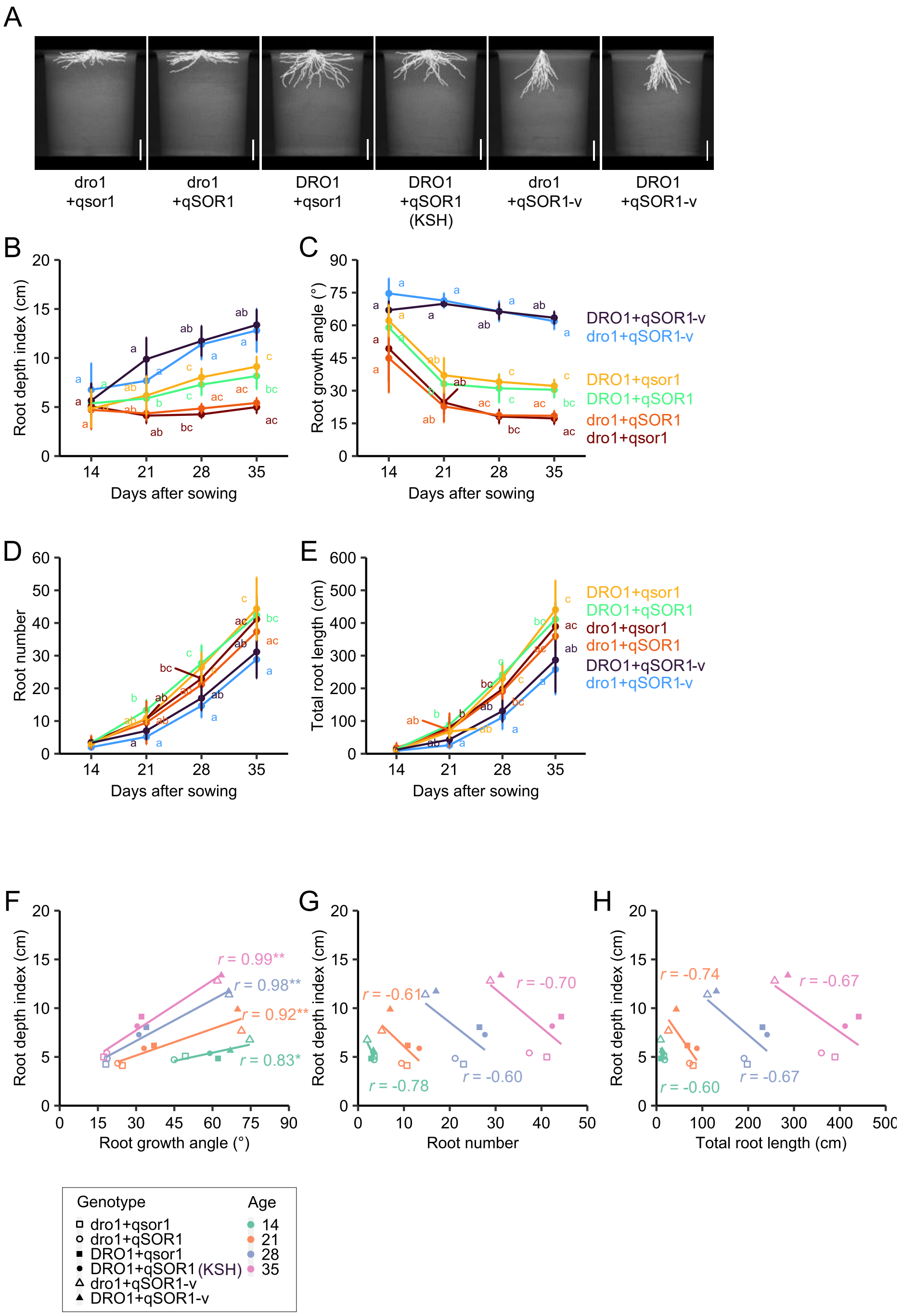


**Figure S9.** RSA analysis of KSH and the five ILs with different *DRO1* and *qSOR1* allele combinations grown in a plant pot under a controlled chamber. (A) Images of RSA in 35-day-old rice plants visualized by X-ray CT and image processing. Scale bars, 5 cm. (B–E) Time-course analyses of (B) root depth index, (C) root growth angle, (D) root number, and (E) total root length. Line charts show mean values ± SD ($n$ = 6 biological replicates). Different lowercase letters indicate significant differences among groups ($p < 0.05$, multiple-comparison Tukey's test). (F–H) Relationships between root depth index and (F) root growth angle, (G) root number, and (H) total root length across the six lines at different plant ages. $r$, Pearson's correlation coefficient (*$p < 0.05$, **$p < 0.01$).

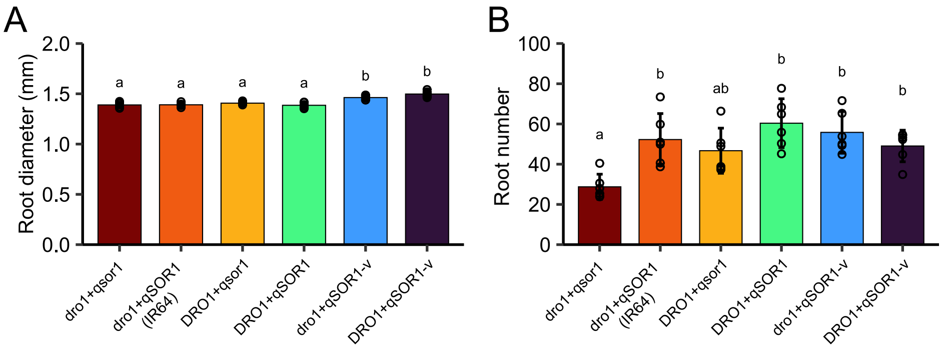


**Figure S10.** Analysis of (A) root diameter and (B) root number in IR64 and the five ILs with different *DRO1* and *qSOR1* allele combinations grown in the paddy field. Bar plots show mean values ± SD ($n$ = 6 biological replicates). Different lowercase letters indicate significant differences among groups ($p < 0.05$, multiple-comparison Tukey's test).

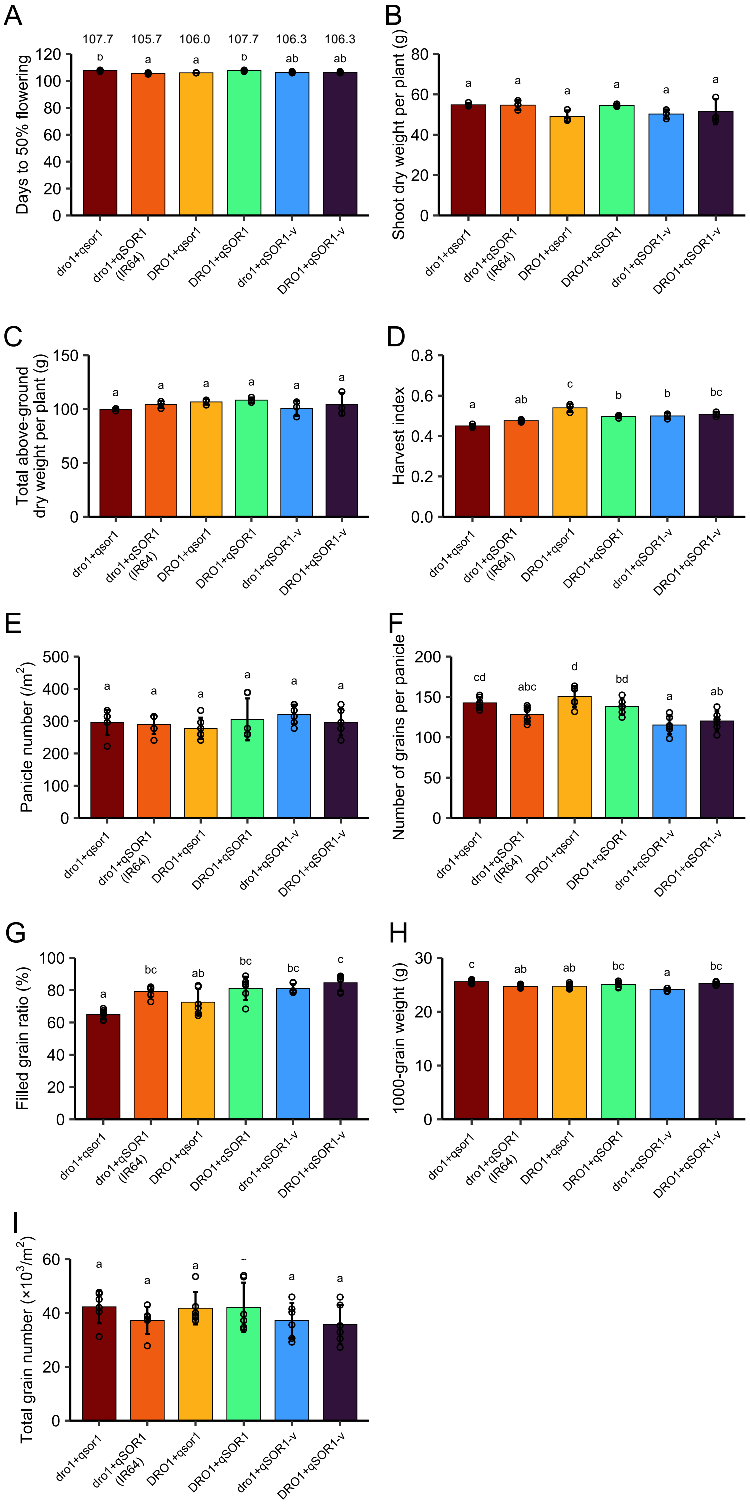


**Figure S11.** Analysis of aboveground traits in IR64 and the five ILs with different *DRO1* and *qSOR1* allele combinations grown in the paddy field. The values represent the days to 50% flowering in (A). (A–D) $n$ = 3 and (E–I) $n$ = 6 biological replicates. Bar plots show mean values ± SD. Different lowercase letters indicate significant differences among groups ($p$ < 0.05, multiple-comparison Tukey's test).

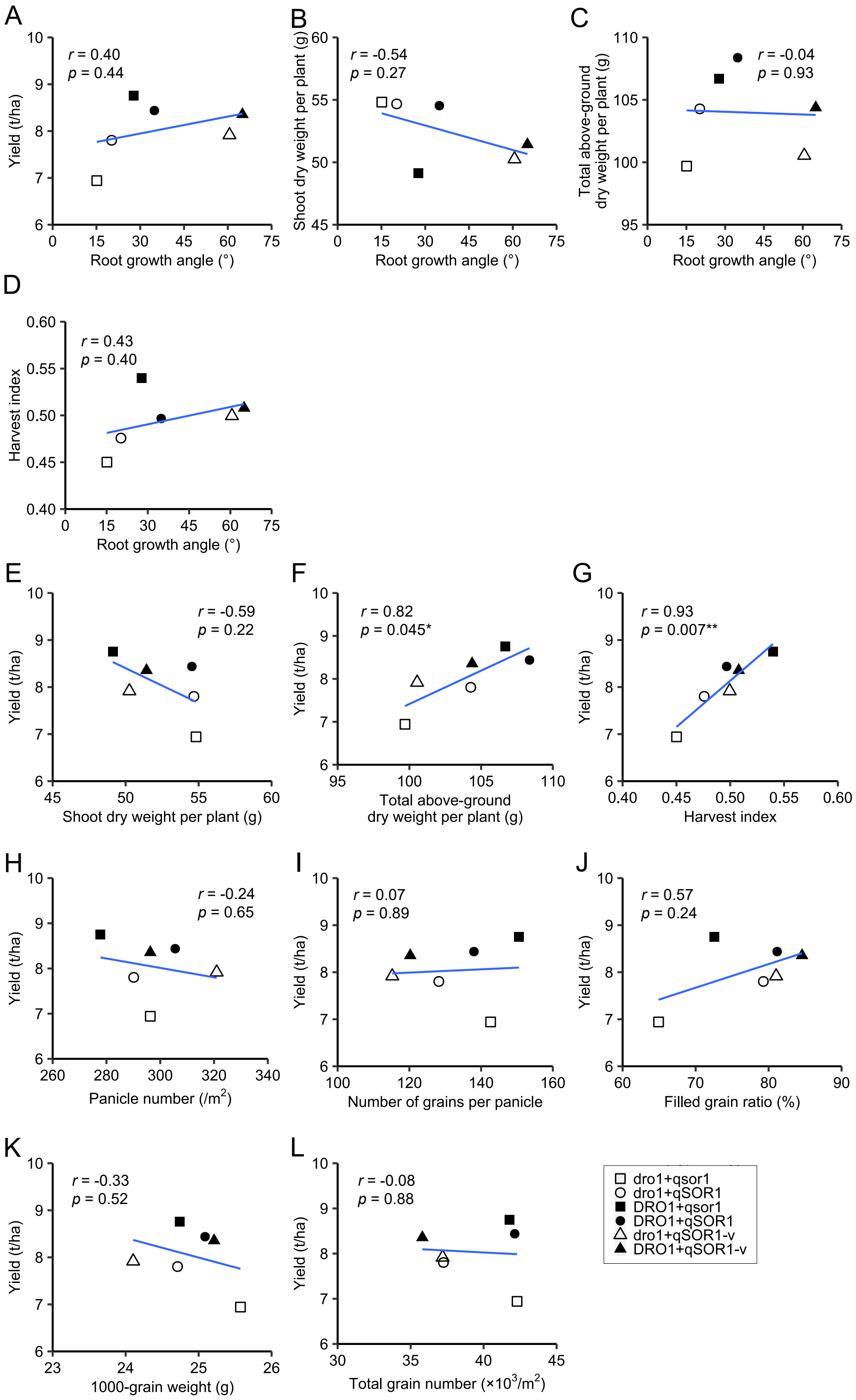


**Figure S12.** Relationships between agronomic traits in IR64 and the five ILs with different *DRO1* and *qSOR1* allele combinations grown in the paddy field. (A–D) Root growth angle and aboveground traits. (E–L) Yield and yield components. *r*, Pearson's correlation coefficient ($*p < 0.05$, $**p < 0.01$).

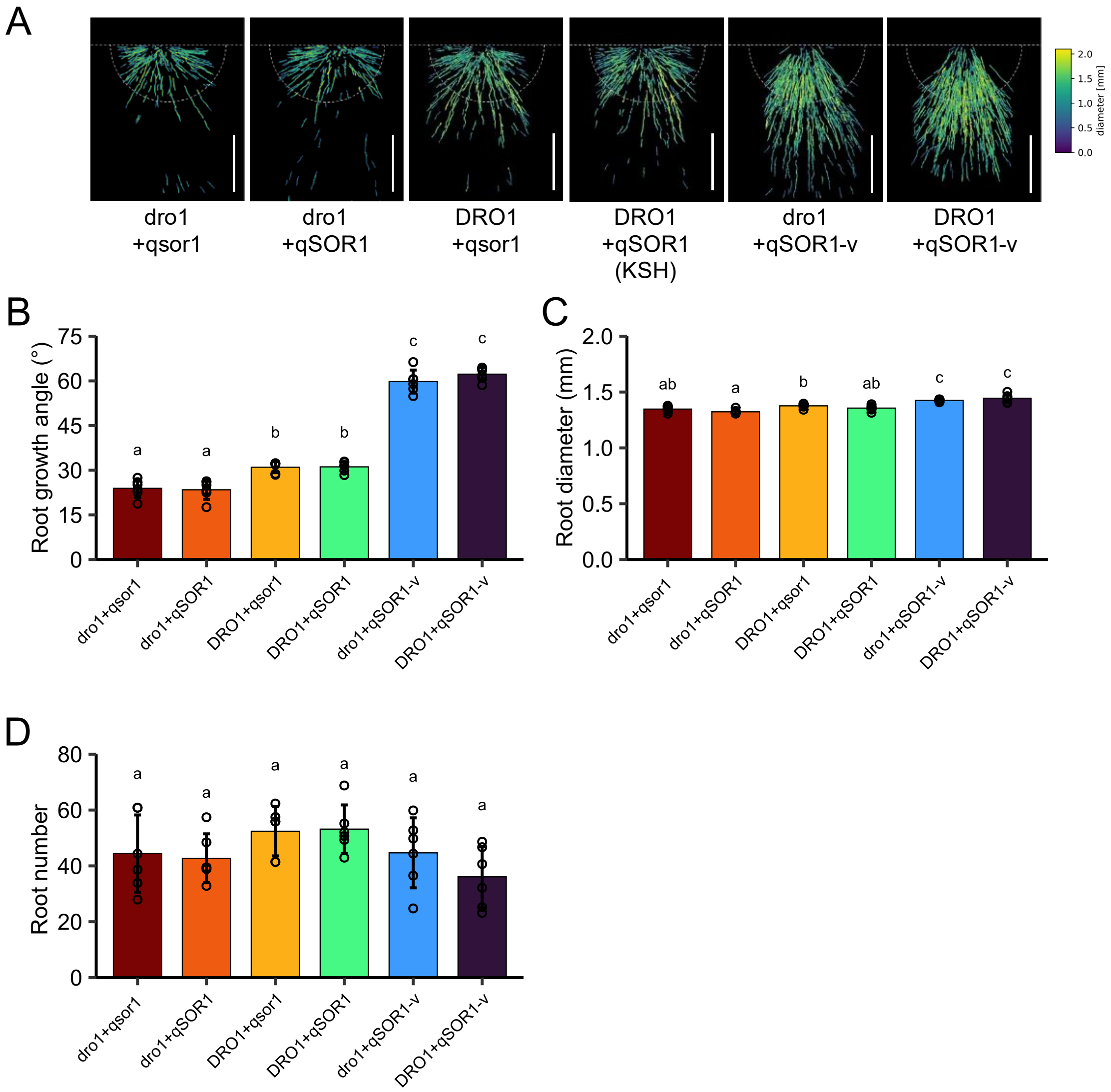


**Figure S13.** Analysis of RSA in KSH and the five ILs with different *DRO1* and *qSOR1* allele combinations grown in the paddy field. (A) Images of the RSA of rice plants at the maturity stage visualized using X-ray CT and image processing. The horizontal dashed lines and half circles indicate the soil surface and the area used for the RSA measurement (radius = 8 cm), respectively. Root color represents root diameter. Scale bars, 5 cm. (B–D) Root growth angle (B), root diameter (C), and root diameter (D) measured in (A). Bar plots show mean values ± SD ($n$ = 6 biological replicates). Different lowercase letters indicate significant differences among groups ($p$ < 0.05, multiple-comparison Tukey's test).

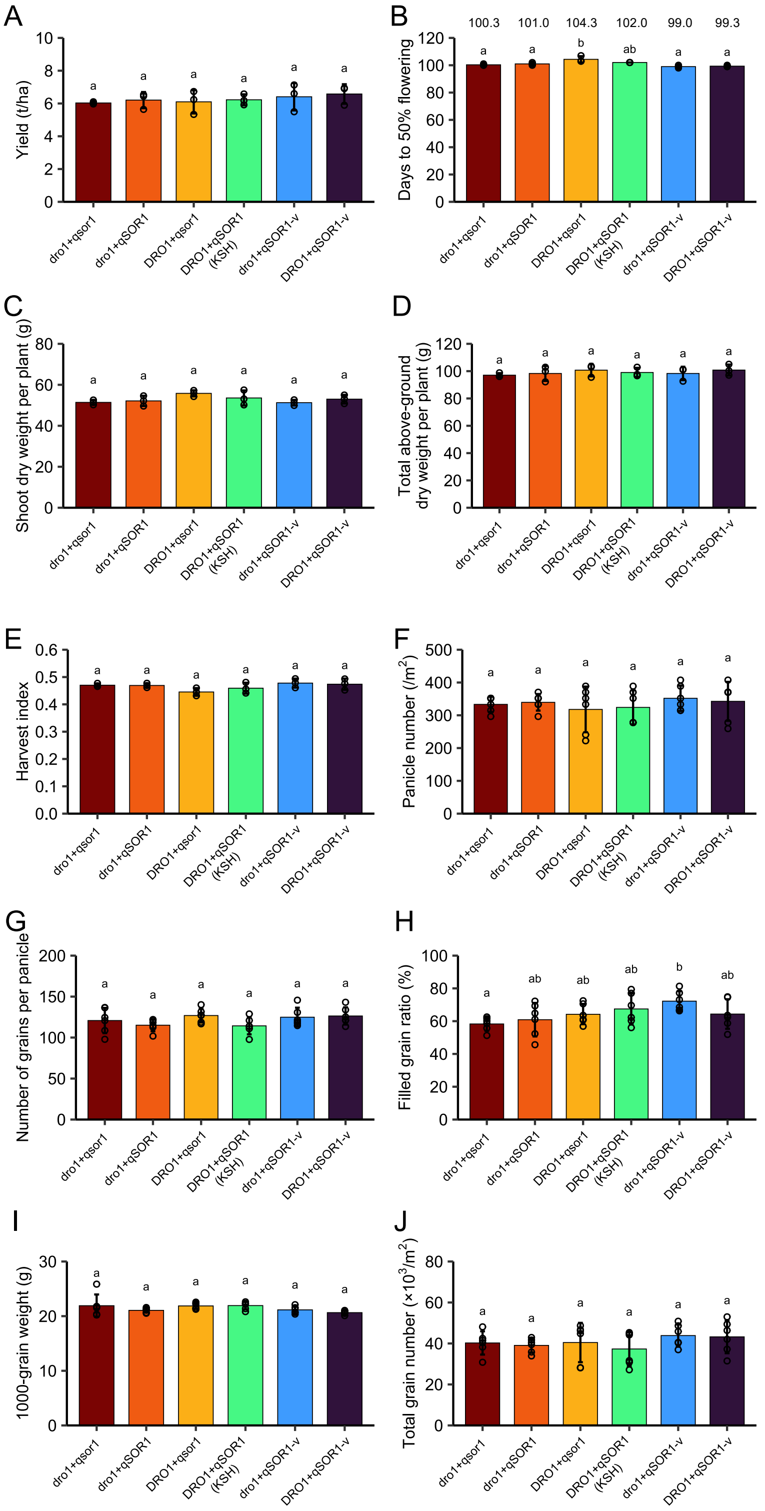


**Figure S14.** Analysis of aboveground traits in KSH and the five ILs with different *DRO1* and *qSOR1* allele combinations grown in the paddy field. The values represent the days to 50% flowering in (B). (A–E) $n$ = 3 and (F–J) $n$ = 6 biological replicates. Bar plots show mean values ± SD. Different lowercase letters indicate significant differences among groups ($p < 0.05$, multiple-comparison Tukey's test).

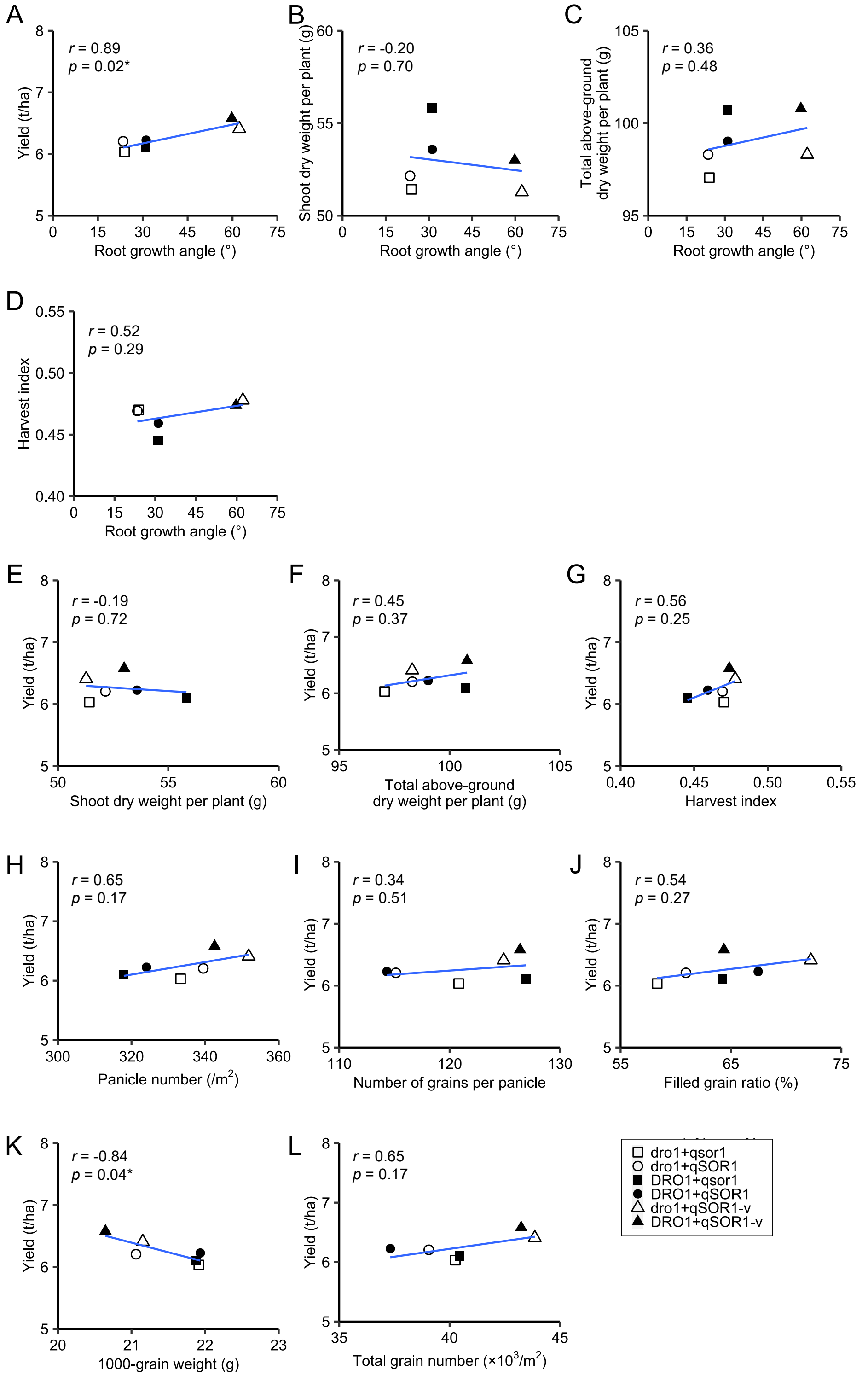


**Figure S15.** Relationships between agronomic traits in KSH and the five ILs with different *DRO1* and *qSOR1* allele combinations grown in the paddy field. (A–D) Root growth angle and aboveground traits. (E–L) Yield and yield components. *r*, Pearson's correlation coefficient (**p* < 0.05, ***p* < 0.01).

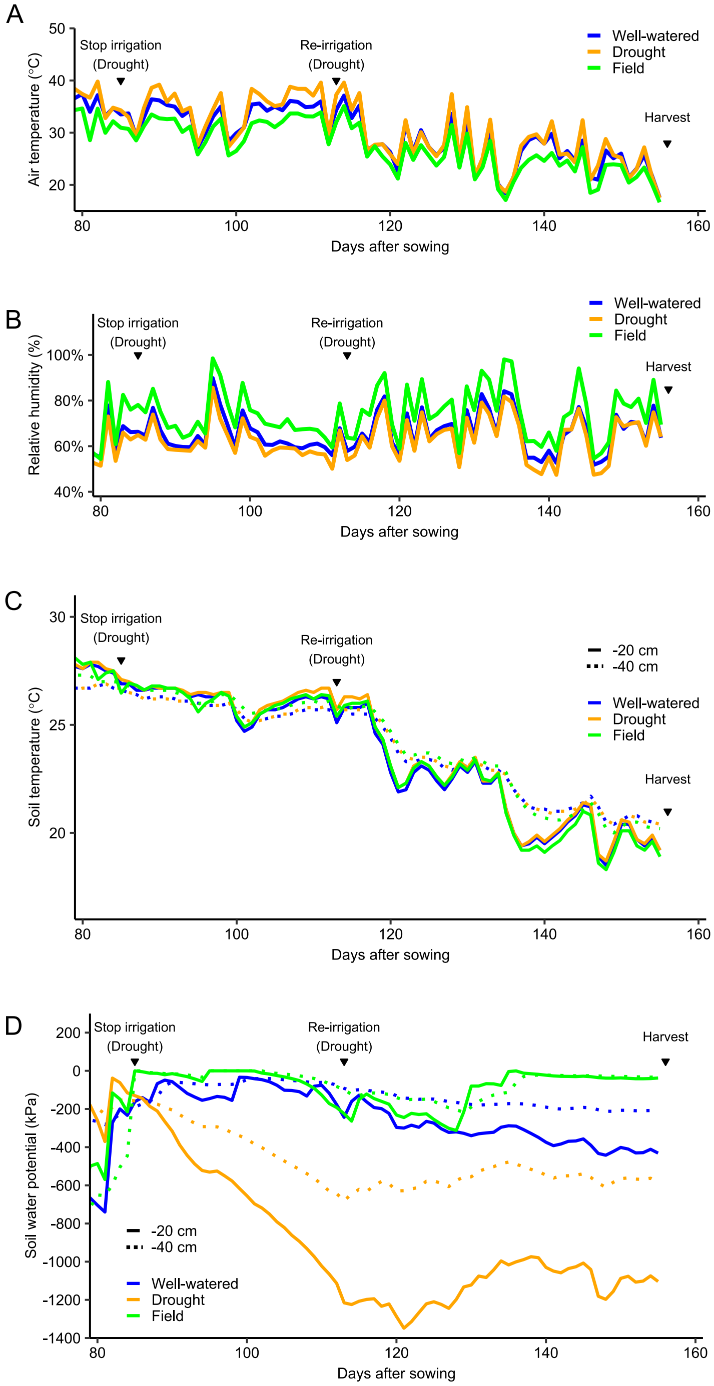


**Figure S16.** Time-course data of (A) air temperature, (B) relative humidity, (C) soil temperature, and (D) soil water potential in the upland field trial. The blue and yellow lines represent data from well-watered and drought stress conditions, respectively, in the greenhouse. The green line represents data from the upland with well-watered conditions for RSA analysis. The inverted triangles indicate the timing of stop irrigation, re-irrigation (for drought treatment), and harvest. The soil parameters were measured at 20 cm (solid lines) and 40 cm (broken lines) below the soil surface in (C) and (D).

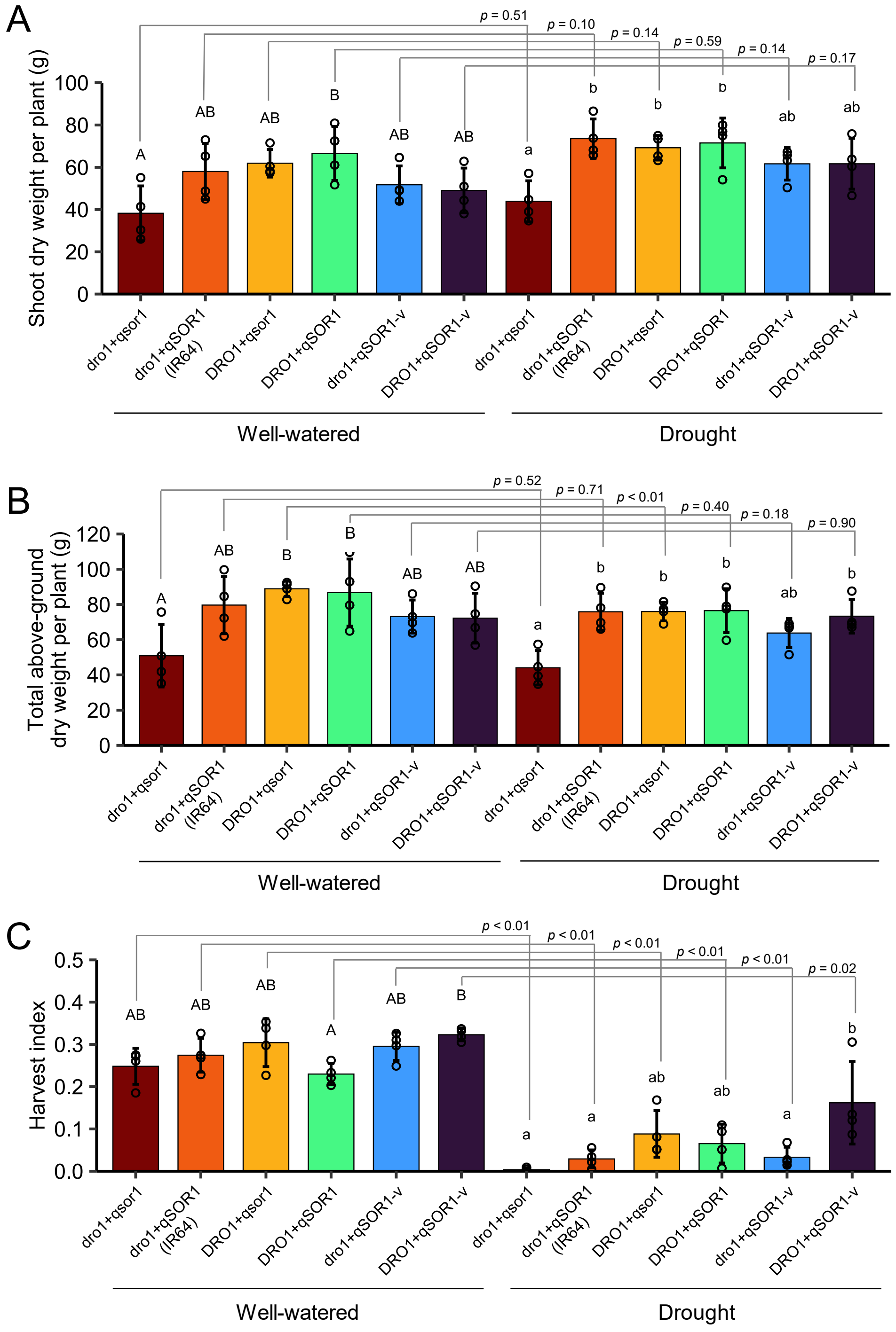


**Figure S17.** Analyses of aboveground traits in IR64 and the five ILs with different *DRO1* and *qSOR1* allele combinations grown in the upland condition. Bar plots show mean values ± SD ($n = 4$ biological replicates). Different letters indicate significant differences among groups ($p < 0.05$, multiple-comparison Tukey's test). $p$ values are derived from a two-tailed Student's $t$-test.

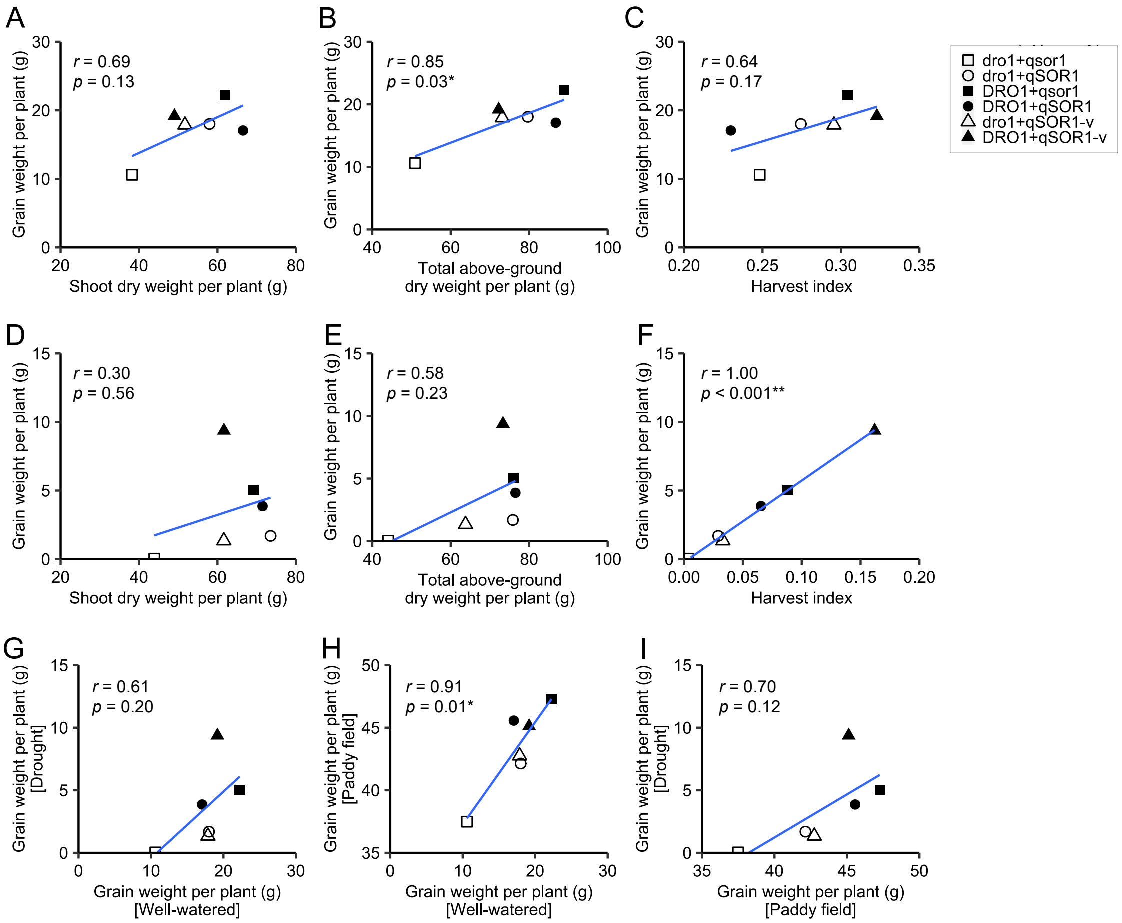


**Figure S18.** Relationships between agronomic traits in IR64 and the five ILs with different *DRO1* and *qSOR1* allele combinations. (A–F) Grain weight per plant and aboveground traits in the upland under (A–C) well-watered and (D–F) drought stress conditions. (G–I) Grain weight per panicle in the different growth conditions. *r*, Pearson's correlation coefficient (**p* < 0.05, ***p* < 0.01).

**Table S1** Description of the 12 rice lines used in this study.

| Allele combination | Line name | Origin of *DRO1* allele | Origin of *qSOR1* allele |
|---|---|---|---|
| (IR64 background) | | | |
| dro1+qsor1 | qsor1_NIL (IR64) | IR64 | GB |
| dro1+qSOR1 | IR64 | IR64 | IR64 |
| DRO1+qsor1 | DRO1+qsor1_PYL (IR64) | KP | GB |
| DRO1+qSOR1 | DRO1_NIL (IR64) | KP | IR64 |
| dro1+qSOR1-v | qSOR1-v_NIL (IR64) | IR64 | 2792M |
| DRO1+qSOR1-v | DRO1+qSOR1-v_PYL (IR64) | KP | 2792M |
| (KSH background) | | | |
| dro1+qsor1 | dro1+qsor1_PYL (KSH) | IR64 | 0909M |
| dro1+qSOR1 | dro1_NIL (KSH) | IR64 | KSH |
| DRO1+qsor1 | qsor1_NIL (KSH) | KSH | 0909M |
| DRO1+qSOR1 | KSH | KSH | KSH |
| dro1+qSOR1-v | dro1+qSOR1-v_PYL (KSH) | IR64 | 2792M |
| DRO1+qSOR1-v | qSOR1-v_NIL (KSH) | KSH | 2792M |

**Table S2** Primers used in this study.

| # | Name | Sequence (5' to 3') | Description |
|---|---|---|---|
| 1 | qSOR1-p1-F1 | TGGATATATTTTGCATGGTTTTTG | PCR of *qSOR1* gDNA fragment for mutant screening |
| 2 | qSOR1-p1-R2 | GAAATGGAGTGAGTAGATGATAACTTG | |
| 3 | SOR-03 | TTCGATTCTTGATTCGACTGG | Sequence of *qSOR1* gDNA fragment for mutant screening |
| 4 | SOR-04U | AGCCAAGTCCACCAAGTCC | |
| 5 | qSOR1Taq02L | CGGCTTATCCTCAAGATAGTATTTCC | |
| 6 | qSOR1-qPCR-Fw | ACGATGCTTCACAAGAGGATGA | Expression analysis ( qRT-PCR) |
| 7 | qSOR1-qPCR-Rv | CGGCTTATCCTCAAGATAGTATTTCC | |
| 8 | OsIAA20-qPCR-Fw | CGGGATTATTTTGTTCACGTTTC | |
| 9 | OsIAA20-qPCR-Rv | CGAGATTTCATTCGTCATGCTTA | |
| 10 | OsUBQ10-qPCR-Fw | GAGCCTCTGTTCGTCAAGTA | |
| 11 | OsUBQ10-qPCR-Rv | ACTCGATGGTCCATTAAACC | |
| 12 | LZY3-qPCR-Fw | CTACACAAGAAAGTTAACAC | |
| 13 | LZY3-qPCR-Rv | CATCGTTACTGCTTCGTC | |
| 14 | ACT2-qPCR-Fw | TCCCTCAGCACATTCCAGCAGATG | |
| 15 | ACT2-qPCR-Rv | AACGATTCCTGGACCTGCCTCATC | |
| 16 | 2792MF_411 | TTCCTTTCGATAGATTCTTGAATTGT | PCR of *LZY3* gDNA fragment with the point mutation (L131F) |
| 17 | 2792MR_374 | ATCTATCGAAAGGAAGATTCGCTAAT | |
| 18 | dqSOR1F_439 | GCTGCCGTTCGACAGGTTCCTCAACT | PCR of *qSOR1* gDNA flagment with the point mutation (L141F) |
| 19 | dqSOR1R_402 | CTGTCGAACGGCAGCAGCAGCTGGCC | |

**Table S3** Operating parameters of the mass spectrometer for IAA analysis.

| | Parameter | Value |
|---|---|---|
| Interface setting | Ion Source | HESI |
| | Probe Heater Temperature | 350° C |
| | Capillary Temperature | 360° C |
| | Sheath gas | 50 arb. |
| | Aux Gas | 5 arb. |
| | S-Lens RF Level | 45% |
| Global setting | Lock mass | *m/z* 130.15903 (+) |
| | Chrom. peak width (FWHM) | 6 sec |
| Experiment setting | Mode | tSIM |
| | Resolution | 70,000 |
| | Isolation window | 1.5 Da |
| | AGC target | 5.00E+04 |
| | Maximum IT | 100 ms |
| | Loop count | 1 |
| | MSX count | 1 |
| | Microscans | 1 |
| | Spectrum data type | Profile |
| | Inclusion list | m/z 176.07060 (+); IAA |
| | | m/z 181.10199 (+); IAA-$d_5$ |
